# TRIM67 drives tumorigenesis in oligodendrogliomas through Rho GTPase-dependent membrane blebbing

**DOI:** 10.1101/2021.11.05.467405

**Authors:** Engin Demirdizen, Ruslan Al-Ali, Ashwin Narayanan, Xueyuan Sun, Julianna Patricia Varga, Bianca Steffl, Manuela Brom, Damir Krunic, Claudia Schmidt, Gabriele Schmidt, Felix Bestvater, Julian Taranda, Şevin Turcan

## Abstract

Oligodendrogliomas are a subtype of isocitrate dehydrogenase (IDH) mutant gliomas defined by the co-deletion of chromosome arms 1p and 19q. Although the somatic genomic alterations of oligodendrogliomas have been well described, transcriptional changes unique to these tumors are not well studied. Here, we identify Tripartite Motif Containing 67 (TRIM67), an E3 ubiquitin ligase with essential roles during neuronal development, as an oncogene distinctly upregulated in oligodendrogliomas. We characterize the function of TRIM67 using high throughput assays, including RNA sequencing, total lysate-mass spectrometry (MS) and co-immunoprecipitation (IP)-MS using human neural progenitor cells and patient-derived glioma tumorspheres constitutively overexpressing TRIM67. Our high throughput data suggest that TRIM67 overexpression alters the abundance of cytoskeletal proteins, which were validated by functional assays, including immunofluorescence (IF) staining, co-IP and western blotting (WB). Additionally, IF staining results indicate that TRIM67 ectopic expression induces formation of membrane blebs in glioma cells, which could be reverted with the nonmuscle class II myosin inhibitor blebbistatin and selective ROCK inhibitor fasudil. GTP pulldown and WB assays further indicate that Rho GTPase/ROCK2 signaling is altered upon TRIM67 ectopic expression. Phenotypically, TRIM67 expression resulted in higher cell motility in wound healing experiments, reduced cell adherence in adhesion assays, accelerated tumor growth and reduced survival in mouse orthotopic implantation models of an oligodendroglioma-derived patient tumorsphere line. Taken together, our results demonstrate that upregulated TRIM67 induces blebbing-based rounded cell morphology through Rho GTPase/ROCK-mediated signaling thereby contributing to glioma pathogenesis.

**Significance:** We identify *TRIM67* as a novel oncogene in oligodendroglioma that leads to increased cell motility, tumor growth, reduced adhesion, and survival in mice. Our results also show that constitutive TRIM67 expression transforms cell morphology from an adherent to a rounded appearance with membrane blebs. Mechanistic alteration of actin cytoskeleton and Rho GTPase signaling upon TRIM67 upregulation underlies the rounded cell structure and the membrane blebbing phenotype. TRIM67-induced blebbing is specifically regulated by RHOA-RAC1-ROCK2 signaling axis. TRIM67 overexpression also alters pathways associated with cell migration and wound healing in various glioma cell lines and human neural progenitor cells, suggesting a general oncogenic mechanism in gliomas. Overall, our study highlights TRIM67 as a novel player orchestrating cytoskeleton, Rho GTPase signaling and bleb-based cell movement, ultimately causing tumorigenic outcomes in oligodendrogliomas.

## Introduction

Lower grade gliomas (LGGs) harbor recurrent mutations in the isocitrate dehydrogenase (IDH) gene. IDH mutant gliomas are classified based on the presence (IDHmut-codel) or absence (IDHmut-noncodel) of chromosome 1p and 19q co-deletions (1, 2). IDHmut-codel and IDHmut-noncodel gliomas differ in their mutational landscape and clinical course (1). IDHmut-codel gliomas, otherwise known as oligodendrogliomas, are slow-growing tumors but can progress to largely incurable high-grade disease. Treatment options for progressive disease are limited, therefore there is a need to develop more effective therapeutics for glioma patients.

While the mutational landscape of IDH mutant gliomas is largely elucidated, the subtype-specific non-mutational mechanisms and their impact on glioma growth are not well understood. In this study, we identify *TRIM67* as a distinctly upregulated gene in IDHmut-codel gliomas. TRIM67 is a member of the large family of tripartite motif (TRIM) containing RING finger E3 ubiquitin ligases (3). TRIM67 is highly expressed in the cerebellum and has been shown to interact with TRIM9 and the netrin receptor DCC and to play a role in neurodevelopment in the mouse brain (4, 5). Further evidence suggests that it antagonizes TRIM9-dependent degradation of VASP, positively regulating filopodia extension and axon branching (6). Most recently, TRIM67 and TRIM9 have been suggested to associate with cytoskeletal proteins and synaptic regulators thus playing a role in exocytosis and endocytosis (7, 8). Therefore, the recently discovered ubiquitin players of neurogenesis, including TRIM67 and TRIM9, may play a role in neuronal development, degeneration, or tumorigenesis (5, 9).

TRIM67 has been reported to play a role in various types of cancer. TRIM67 has been found to downregulate Ras signaling and activate differentiation in neuroblastoma cells (10). It has been shown to activate the p53 pathway and suppress colorectal cancer in mice (11). Another study indicated that TRIM67 promotes NF-kB signaling and apoptosis in lung cancer cells (12). In non-small cell lung cancer, TRIM67 upregulates Notch signaling pathway and causes proliferation, migration, and invasion (13). Although the role of TRIM67 in normal neuronal development has been elegantly shown, it is unclear whether TRIM67 has an oncogenic role in gliomas.

The aim of this study is to characterize the mechanistic role of TRIM67 in glioma pathogenesis. To this end, we used human neural progenitor cells and patient-derived glioma tumorsphere lines with or without IDH mutation and 1p/19q co-deletion to investigate the function of TRIM67 in gliomas. We find that TRIM67 induces membrane blebbing through Rho GTPase-mediated signaling, leading to reduced adhesion, and increased cell motility and tumor expansion.

## Results

### *TRIM67* is overexpressed in IDHmut-codel gliomas

To determine transcriptional differences between IDHmut-codel and IDHmut-noncodel gliomas, we analyzed the LGG samples from The Cancer Genome Atlas (TCGA) RNA sequencing (RNA-seq) dataset. We identified *TRIM67* as one of the top significantly upregulated genes in IDHmut-codel tumors (Figure 1A). TRIM67 exhibited approximately 3.5-fold increased expression compared with IDHmut-noncodel or IDH wild-type LGGs (Figure 1B). Our results indicate that TRIM67 may be overexpressed through an epigenetic control mechanism that includes promoter hypomethylation and open chromatin structure (Figure 1C-D). A pan-cancer comparison showed that *TRIM67* expression is higher in IDHmut-codel gliomas than in most other tumor types, including glioblastomas (GBM), with a few exceptions such as neuroblastoma (NBL) and adrenal gland tumors (PCPG) (Supplemental Fig. 1A). This suggests that TRIM67 may play a role in the pathogenesis of oligodendrogliomas.

**Figure 1.**
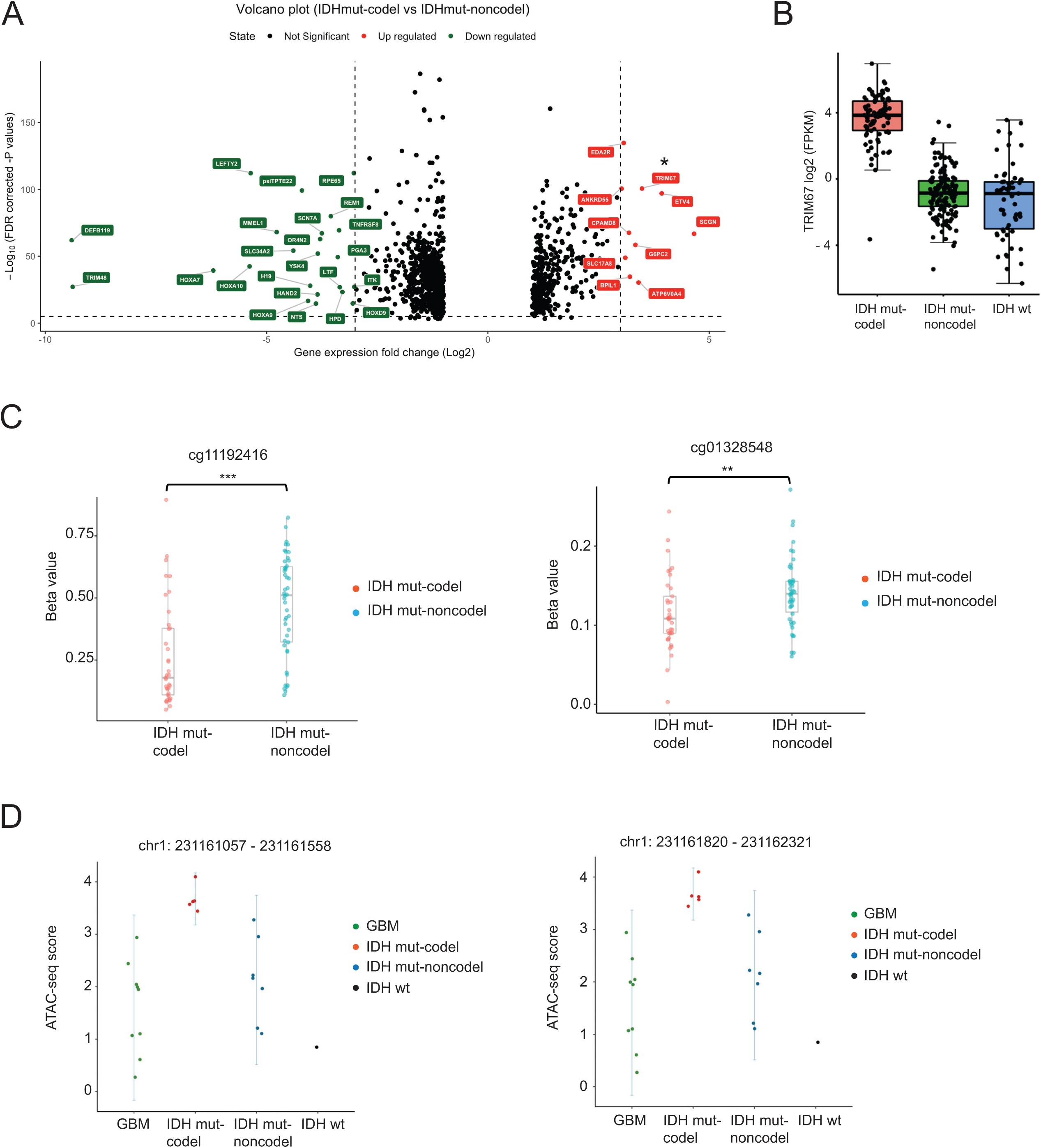
*TRIM67* is upregulated in oligodendrogliomas. **A)** The volcano plot for the altered genes between IDHmut-codel (oligodendroglioma) and IDHmut-noncodel gliomas using TCGA data. The significance threshold for -log_10_(FDR-corrected p values) is 1.4. Thresholds for log_2_ gene expression are -3 and 3. Upregulated genes are depicted in red, downregulated genes are depicted in green and non-altered genes are depicted in black. *TRIM67* is designated by asterisk. **B)** The bar graph comparison of log_2_ expression for *TRIM67* between IDHmut-codel (oligodendroglioma), IDHmut-noncodel and IDH wild type gliomas using TCGA data. **C)** EPIC methylation beta values at two different *TRIM67* promoter regions comparing data from 1p19q_codel and astrocytoma patients. Two-sided t-test was performed, ** p<0.01 and *** p<0.001. **D)** ATAC-seq scores at TRIM67 promoter and 5’ UTR regions comparing different glioma subtypes.

### TRIM67 alters pathways regulating cytoskeletal organization

To uncover the role of TRIM67 in gliomas, we used cultured human neural progenitor cells (hNPCs) and four patient-derived glioma tumorsphere lines, consisting of one IDH-mutant oligodendroglioma line (TS603) and three IDH-wild-type lines (S24, TS600, and TS543). Due to the paucity of IDH mutant oligodendroglioma lines, we used additional IDH-wild-type glioma lines to study TRIM67 function. We engineered all five lines to constitutively overexpress Flag-tagged TRIM67 (Supplemental Fig. 1B). To gain insight into the function of TRIM67, we performed RNA-seq on the TRIM67 overexpressing and control lines and identified significantly altered genes upon TRIM67 overexpression (Supplemental Table 1). In the TS603 line, cytoskeleton-associated genes accounted for the majority of differentially expressed genes (Figure 2A). KEGG pathway and gene ontology analysis indicated that cytoskeleton and focal adhesion were among the most significantly altered pathways (Figure 2B). Comparison of pathways altered by TRIM67 across different cell lines revealed significant alteration of cytoskeleton-related pathways, such as paxillin signaling and actin cytoskeleton signaling (Figure 2C).

**Figure 2.**
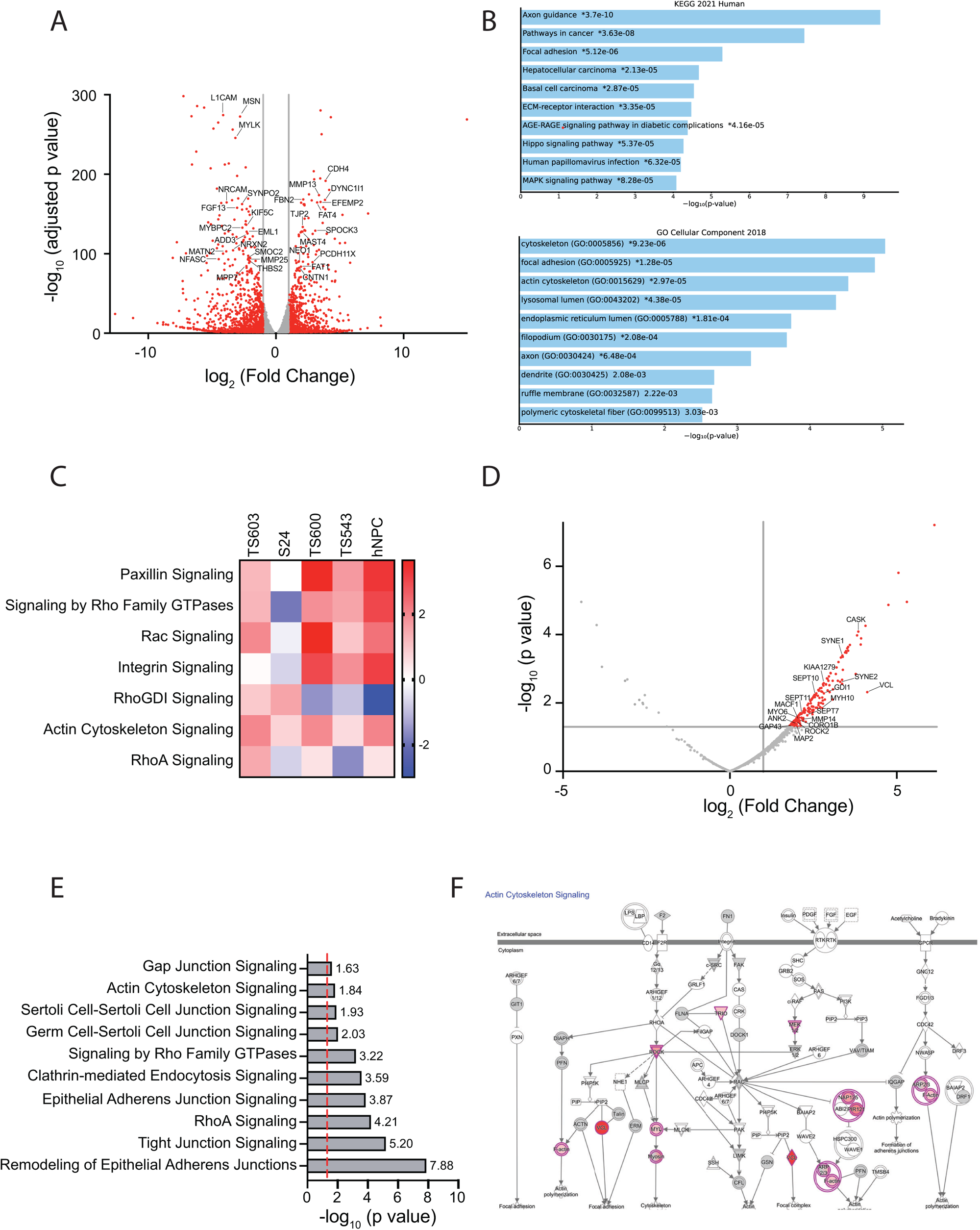
Overexpressed TRIM67 associates with and alters cytoskeleton signaling. **A)** Volcano plot for TS603 RNA-Seq upon TRIM67 overexpression. The genes with -log_10_(padj) value greater than 1.4 are depicted. Thresholds for log_2_ fold change are -1 and 1. The altered genes of interest are marked and shown on the plot. **B)** Top 10 significantly-enriched KEGG pathways and GO cellular components upon TRIM67 overexpression plotted by using EnrichR. C) Heatmap comparison of RNA-Seq pathways for 5 different cell lines using IPA and Prism. D) Volcano plot for Co-IP MS. The significance threshold for -log_10_(padj) was set to 1.4. Threshold for log_2_ fold change is 1. The significant proteins are depicted in red, and non-significant proteins are depicted in gray. The proteins of interest are marked and shown on the plot. **E)** Top 10 significantly-enriched cytoskeletal pathways from Co-IP MS experiment. The significance threshold for -log_10_(p value) was set to 1.4, which is shown by the dashed red line. **F)** The scheme of actin cytoskeleton signaling created by using IPA. The proteins marked in red were detected from Co-IP MS.

To further shed light on TRIM67-dependent mechanisms, we performed mass spectrometry (MS)-based approaches using total cell extracts or TRIM67 pulldown using Flag antibody (Supplemental Table 2). In the TRIM67-Flag co-immunoprecipitation (co-IP)-MS experiment, we observed that a significant proportion of TRIM67-interacting partners contained cytoskeletal proteins (Figure 2D) and actin cytoskeleton pathways were significantly enriched (Figure 2E). Notably, several members of the actin cytoskeleton pathway were detected to directly or indirectly interact with TRIM67 (Figure 2F). Similarly, the total cell extract-MS approach using two different TRIM67 overexpressing cell lines (TS603, S24) also showed a significant change in the abundance of cytoskeletal proteins (Supplemental Fig. 1C-D). Pathway analysis indicated that cytoskeletal and cell adhesion pathways were significantly altered among the differentially enriched proteins (Supplemental Fig. 1E-F). Detailed analysis of protein abundance of the members of the actin cytoskeleton pathway revealed enrichment of these proteins in TRIM67 overexpressing cell lines (Supplemental Fig. 1G-H). To this end, our high-throughput data suggest that TRIM67 may lead to alterations in the cellular cytoskeleton and focal adhesion.

### TRIM67 partially interacts with cytoskeletal proteins and changes their structure or expression

Next, we aimed to address how TRIM67 regulates cytoskeletal proteins. To this end, we performed immunofluorescence (IF) staining for a set of cytoskeletal proteins, including actin, tubulin, F-actin and MAP1B. Using transiently transfected HEK cells, we observed that actin-RFP and tubulin-RFP exhibited aggregate-like structures upon TRIM67 overexpression (Figure 3A), pointing to the changes in the cytoskeletal structure. Interestingly, F-actin levels decreased significantly upon TRIM67 overexpression (Figure 3B), by approximately 4-fold in hNPC and 2-fold in TS600 (Figure 3C). To test whether TRIM67 associated with structural proteins, we applied a co-staining approach. Although ectopically expressed TRIM67 was localized within the cytoplasm, we did not observe a complete overlap with actin, tubulin and MAP1B. (Supplemental Fig. 2A-C). Notably, we detected a substantial increase in MAP1B protein levels in TRIM67 overexpressing TS603 cells (Supplemental Fig. 2D). Taken together, these results suggest that TRIM67 alters the distribution of several cytoskeleton-associated proteins.

**Figure 3.**
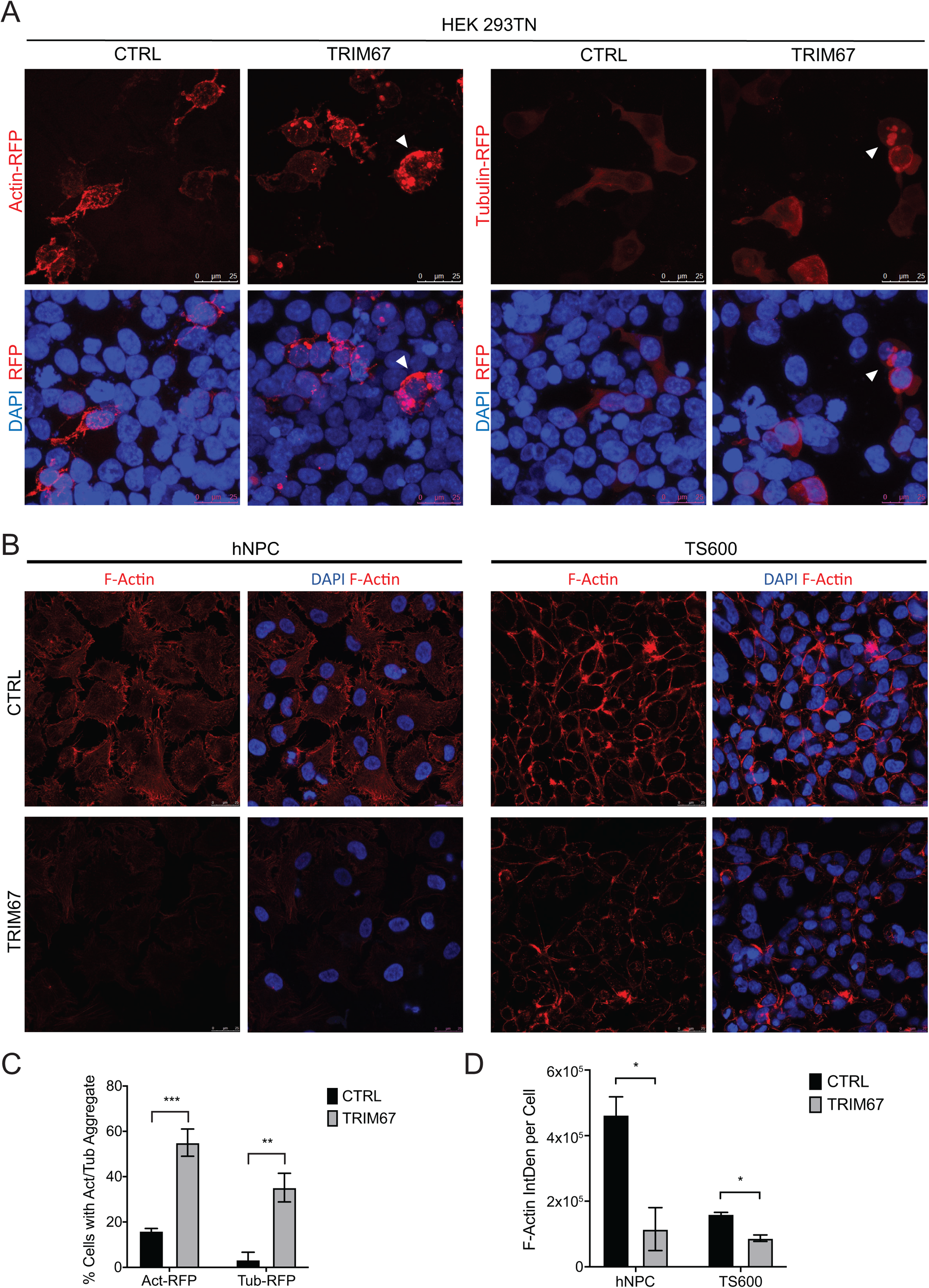
TRIM67 overexpression alters cytoskeletal proteins. **A)** Representative IF images of HEK293TN cells upon control and TRIM67 overexpression conditions. Actin-RFP staining is shown on the left panel, Tubulin-RFP staining is shown on the right panel. RFP in red and DAPI in blue. The cells of interest are marked by white triangles. Scale bar indicates 25 μm. **B)** Representative IF images of cells upon control and TRIM67 overexpression conditions. hNPC cells are shown on the left panel, TS600 cells are shown on the right panel. F-Actin in red and DAPI in blue. Scale bar indicates 25 μm. **C)** Quantification for part A, illustrating the percentage of cells with actin or tubulin aggregates upon control and TRIM67 overexpression conditions. Error bars indicate the standard error of the mean. Two-sided t-test was performed, ** p<0.01 and *** p<0.001. **D)** Quantification for part B, illustrating the F-Actin integrated density per cell in hNPC and TS600 cells upon control and TRIM67 overexpression conditions. Error bars indicate the standard error of the mean. Two-sided t-test was performed, * p<0.05.

### Ectopic TRIM67 expression induces the formation of membrane blebs

The next question we investigated was how TRIM67 alters the cytoskeleton. Confocal images of F-actin staining in HEK cells transiently transfected with TRIM67 or empty vector showed that the morphology of epithelial-like adherent cells changed to a round cell structure compared to control cells which exhibit a less-adherent and spreading phenotype (Figure 4A). Remarkably, both differential interference contrast (DIC) and fluorescent F-actin images showed that transgenic TRIM67 expression induces membrane blebs and TRIM67 clusters adjacently to the blebs (Figure 4A).

**Figure 4.**
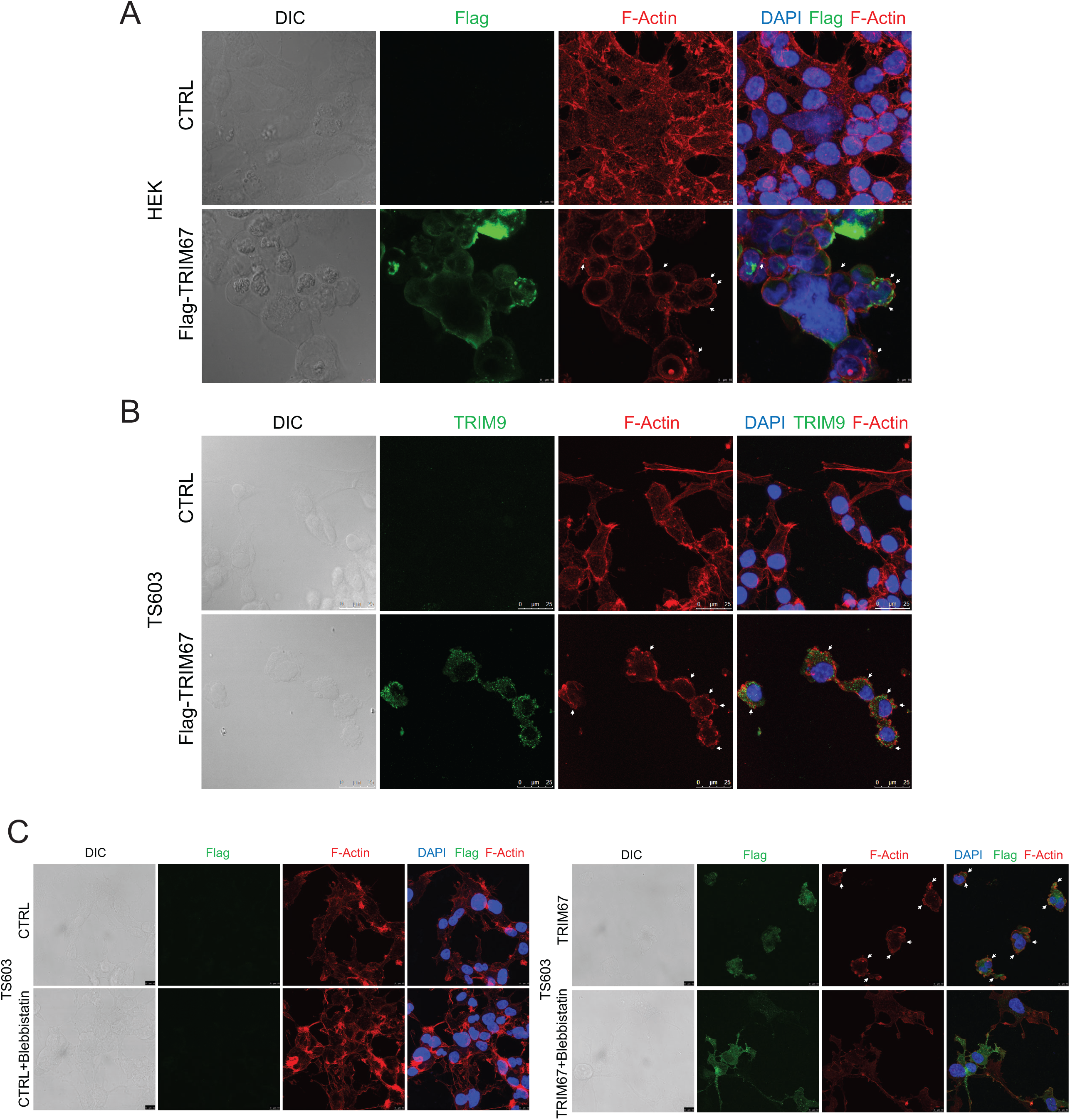
TRIM67 overexpression induces membrane blebbing and rounded cell morphology. **A)** Representative IF images of HEK293-TN cells upon control and TRIM67 overexpression conditions. DIC indicates bright field images, Flag in green, F-Actin in red and DAPI in blue. Membrane blebs are marked by white arrows. Scale bar indicates 10 μm. **B)** Representative IF images of TS603 cells upon control and TRIM67 overexpression conditions. DIC indicates bright field images, TRIM9 in green, F-Actin in red and DAPI in blue. Membrane blebs are marked by white arrows. Scale bar indicates 25 μm. **C)** Representative IF images of TS603 cells with and without blebbistatin treatment upon control and TRIM67 overexpression conditions. Control cells are shown in the left panel, TRIM67-overexpressing cells are shown in the right panel. DIC indicates bright field images, Flag in green, F-Actin in red and DAPI in blue. Membrane blebs are marked by white arrows. Scale bar indicates 10 μm.

Next, we tested the interaction of TRIM67 with TRIM9, the TRIM family member with the closest sequence homology to TRIM67. Both proteins were previously found to associate with cytoskeleton (8), regulate filopodia extensions (6) and play essential roles in axon branching and synaptic vesicle formation (7). By western blotting, we detected a substantial increase in TRIM9 protein levels in three different cell lines (Supplemental Fig. 3A-B). We also confirmed the physical interaction and colocalization of TRIM67 and TRIM9 by co-IP and IF staining (Supplemental Fig. 3C-F). Remarkably, in HEK cells, transient TRIM9-GFP localized to cytoskeletal mesh-like fibers (Supplemental Fig. 3E). Upon TRIM67 co-expression, this specific localization pattern was converted into a vesicle-like aggregate structure, in which TRIM67 and TRIM9 colocalize (Supplemental Fig. 3E). The change in TRIM9 localization upon co-expression of TRIM67 was validated by using Myc-TRIM9 construct (Supplemental Fig. 3F), indicating that the altered localization is not due to an artifact caused by the GFP-tag. Endogenous TRIM9 was also detected to cluster with F-actin in TRIM67-induced membrane blebs in the TS603 oligodendroglioma cell line (Figure 4B). To further prove the role of TRIM67 in membrane blebbing, we used the nonmuscle class II myosin inhibitor blebbistatin to inhibit the formation of blebs. Notably, blebbistatin had no appreciable effect on the F-actin structure and cytoskeleton morphology of control cells (Figure 4C). Interestingly, TRIM67-induced membrane bleb formation was substantially inhibited and reverted to a more adherent phenotype with longer protrusions in the TRIM67 overexpressing cells under blebbistatin treatment (Figure 4C). Overall, these data suggest that TRIM67 ectopic expression triggers membrane blebbing and interacts with and alters the cytoskeletal proteins.

### TRIM67-driven membrane blebbing is linked to Rho GTPase signaling

Next, we addressed the mechanism by which TRIM67 induces membrane blebbing. To do this, we compared our data from independent MS experiments, including total cell extract-MS and co-IP-MS, and analyzed the overlap of differentially active pathways. We found that the actin cytoskeleton signaling and signaling by Rho GTPase pathways were among the top hits (Figure 5A). Similarly, in the RNA-seq datasets, the Rho GTPase-related pathways were also differentially enriched (Figure 2C). Rho GTPase signaling is known to play major roles in cytoskeleton dynamics and membrane blebbing mechanism (14). This led us to hypothesize that TRIM67 might regulate Rho GTPase signaling to stimulate membrane blebbing. Indeed, detailed examination of those pathways using Ingenuity Pathway Analysis (IPA) tool indicated that several members of Rho GTPase pathways were altered, including RHOA and RAC signaling (Figure 5A-C). We further performed a GTP pulldown assay to detect the activity of GTP-bound Rho GTPases. This implicated an enhanced RHOA activity but reduced RAC1 activity in oligodendroglioma cell line upon TRIM67 overexpression (Figure 5D). Consistently, RHOA-ROCK axis is reported as an inducer, while RAC1 is reported as a suppressor of membrane blebbing(15). Indeed, we also detected a substantial increase in ROCK2 protein levels upon TRIM67 overexpression (Figure 5E). To further test the involvement of ROCK signaling in TRIM67-driven blebbing, we used selective ROCK inhibitor Fasudil (HA-1077). Of note, Fasudil treatment does not appear to influence F-actin cytoskeletal structure in control cells (Figure 5F). Intriguingly, TRIM67-driven membrane blebbing was prominently reverted under Fasudil treatment (Figure 5F). These results suggest that TRIM67-induced membrane blebbing is due to altered Rho GTPase/ROCK signaling.

**Figure 5.**
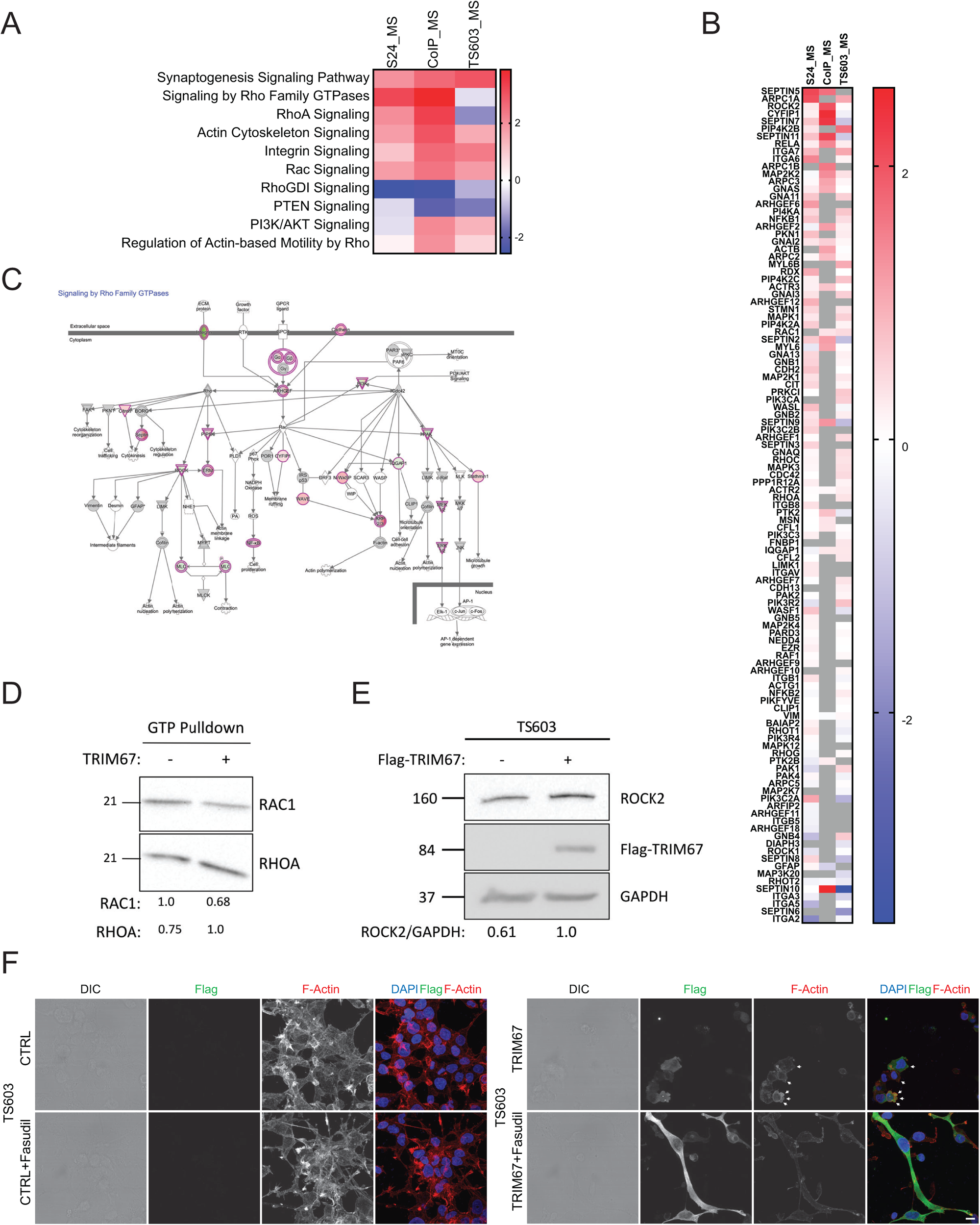
TRIM67 induces membrane blebbing through Rho GTPase/ROCK signaling. **A)** Heatmap comparison of pathways for S24_MS, Co-IP_MS and TS603_MS experiments using IPA and Prism. **B)** Heatmap comparison of signaling by Rho family GTPases pathway for S24_MS, Co-IP_MS and TS603_MS experiments using IPA and Prism. **C)** The scheme of signaling by Rho family GTPases created by using IPA. The proteins marked in red are upregulated, whereas those marked in green are downregulated in S24_MS experiment. **D)** Immunoblotting for RAC1 and RHOA upon GTP pulldown under control and TRIM67 overexpression conditions in TS603 cells. The normalized quantification for RHOA and RAC1 is shown below. **E)** Immunoblotting for Flag-TRIM67, ROCK2 and GAPDH under control and TRIM67 overexpression conditions in TS603 cells. GAPDH serves as a loading control. The normalized quantification for ROCK2 is shown below. **F)** Representative IF images of TS603 cells with and without fasudil treatment upon control and TRIM67 overexpression conditions. Control cells are shown in the left panel, TRIM67-overexpressing cells are shown in the right panel. DIC indicates bright field images, Flag in green, F-Actin in red and DAPI in blue. Membrane blebs are marked by white arrows. Scale bar indicates 10 μm.

### TRIM67 expression promotes cell migration

Membrane blebbing could cause increased cell migration or apoptosis (16).To rule out whether TRIM67 plays a role in apoptosis induction, we performed IF staining for apoptotic markers, including cleaved Caspase3 and cleaved PARP. Despite the presence of membrane blebbing, we did not detect apoptosis in glioma cell lines TS600 and TS603 upon TRIM67 overexpression (Supplemental Fig. 4A-B). Then to gain insights into how cell migration and invasion are influenced by TRIM67 expression, we performed western blots to test the expression of several markers involved in migration and invasion. We observed a significant upregulation in phospho-FAK, phospho-Akt and vimentin but a significant reduction in tensin-2 levels upon TRIM67 overexpression using different glioma cell lines (Figure 6A-B). Tensin-2 has been shown to negatively regulate Akt/PKB signaling pathway and consequently cell proliferation and migration(17). This suggests that ectopic TRIM67 expression could potentially activate cell migration and invasion pathways. To further test this hypothesis, we performed wound healing assays using a neuroblastoma cell line with endogenous TRIM67 expression (SK.N.BE2), and a TRIM67-deficient glioblastoma cell line (TS600) (Figure 6C-D, Supplemental Fig. 5A). SK.N.BE2 cells exhibited significantly reduced migration upon TRIM67 shRNA knockdown (Figure 6C, Supplemental Fig. 5A). Similarly, TS600s had increased cell migration capacity upon TRIM67 overexpression (Figure 6D, Supplemental Fig. 5A). Concomitantly, we further observed a significantly reduced surface adhesiveness of TRIM67-overexpressing TS603 cells on all five distinct coating matrices (Figure 6E-F). These results indicate that TRIM67 leads to enhanced cell migration with poor cell adhesion, which is a hallmark of cancer.

**Figure 6.**
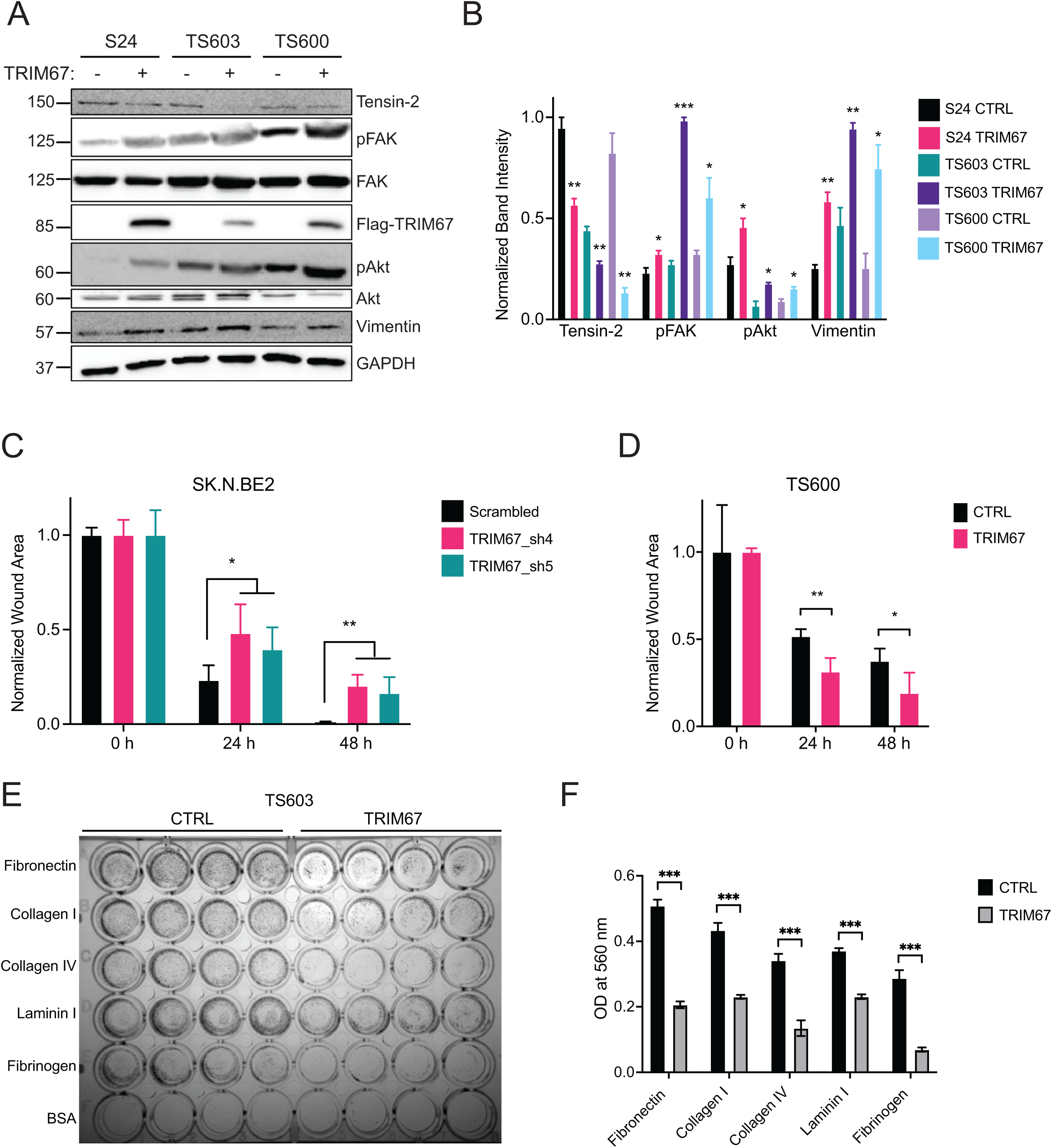
TRIM67 expression induces cell migration. **A)** Immunoblotting for Tensin-2, pFAK, FAK, Flag-TRIM67, pAkt, Akt and GAPDH under control and TRIM67 overexpression conditions in S24, TS603 and TS600 cells. GAPDH serves as a loading control. **B)** The normalized quantification of part A for Tensin-2, pFAK, pAkt and Vimentin. pFAK and pAkt were further normalized to FAK and Akt, respectively. Error bars indicate the standard error of the mean. Two-sided t-test was performed, * p<0.05, ** p<0.01 and *** p<0.001. **C)** The quantification of Supplemental Fig 5A for normalized wound area in SK.N.BE2 cells upon treatments. Error bars indicate the standard error of the mean. Two-sided t-test was performed, * p<0.05 and ** p<0.01. **D)** The quantification of Supplemental Fig 5A for normalized wound area in TS600 cells upon treatments. Error bars indicate the standard error of the mean. Two-sided t-test was performed, * p<0.05 and ** p<0.01. **E)** Cell adhesion assay showing control and TRIM67-overexpressing TS603 cells on five different coating matrices. BSA serves as negative control. **F)** The quantification of part E for normalized OD at 560 nm. Error bars indicate the standard error of the mean. Two-sided t-test was performed, *** p<0.001.

### TRIM67 overexpression leads to increased tumor growth and reduced survival *in vivo*

To address whether TRIM67 is oncogenic *in vivo*, we orthotopically implanted empty vector or TRIM67 overexpressing TS603 glioma cells into the frontal lobes of nude mice. We observed significantly increased tumor growth in mice injected with TRIM67-overexpressing cells by bioluminescence imaging (Supplemental Fig 5B). Quantification of mean flux for individuals and subgroups also indicated an accelerated tumor growth upon TRIM67 overexpression, reaching to significance on day 18 post-implantation (Figure 7A-C). To further analyze the tumors, we dissected the brains from control and TRIM67 groups in triplicates and performed H&E, Flag and Ki-67 staining at two selected brain coronal regions. This experiment confirmed sustained constitutive expression of TRIM67 *in vivo* and showed a significantly higher tumor burden associated with TRIM67 overexpression as measured by H&E and deformation in brain anatomy (Figure 7E). Furthermore, TRIM67 overexpression in mice led to a significant decrease in survival (Figure 7D). Taken together, TRIM67-expressing cells in the mice brains resulted in advanced tumor growth with poor survival.

**Figure 7.**
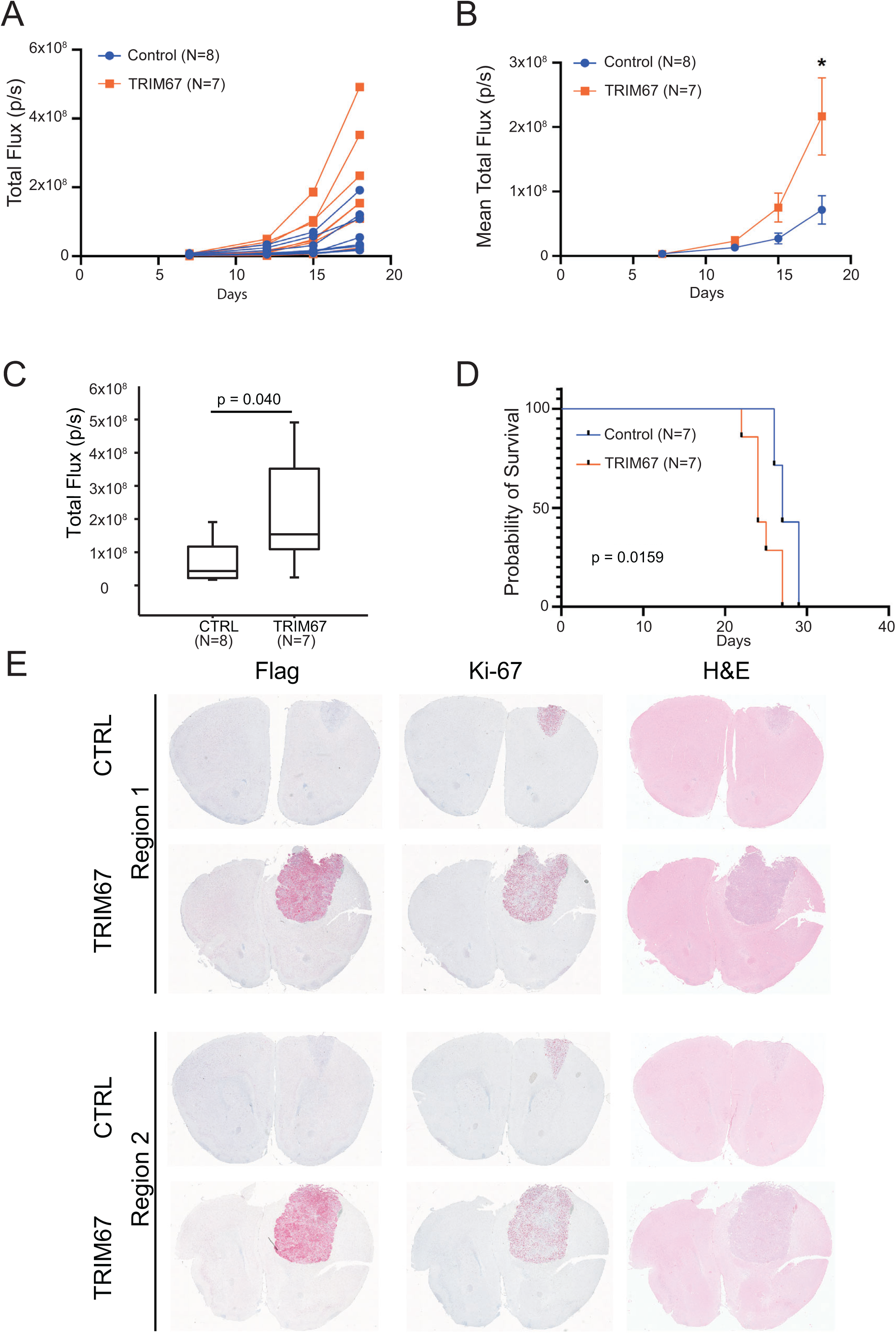
TRIM67 overexpression leads to advanced tumor growth and poor survival in mice. **A)** The graph of the individual tumor growth for each mouse in control (n=8) and TRIM67 (n=7) groups on day 7, 12, 15 and 18. **B)** The graph of the average tumor growth in control (n=8) and TRIM67 (n=7) groups on day 7, 12, 15 and 18. Error bars indicate the standard error of the mean. Two-sided t-test was performed, * p<0.05. **C)** The boxplot of the average tumor growth on day 18 in control (n=8) and TRIM67 (n=7) groups. Error bars indicate the standard error of the mean. Two-sided t-test was performed, p=0.040. **D)** Kaplan Meier curve for the survival analysis of control (n=7) and TRIM67 (n=7) groups. Log rank test was performed, p=0.0159. **E)** IHC staining images for Flag, Ki-67 and H&E using brain sections at two different regions from representative control and TRIM67 animals.

In conclusion, our study identified *TRIM67* as a novel oncogene in oligodendrogliomas. Mechanistically, our results show that overexpressed TRIM67 alters Rho GTPase signaling and induces membrane blebbing, thereby activating the tumor migration and expansion with higher mortality.

## Discussion

Here we identify that TRIM67 is distinctly upregulated in oligodendrogliomas. Our results indicate that TRIM67 acts as an oncogene through the activation of Rho GTPase signaling-dependent membrane blebbing mechanism, leading to cell motility and tumor progression. Our findings illustrate a crucial oncogenic role of TRIM67 in gliomas apart from its essential biological roles in neural development, axon guidance, filopodia dynamics, exocytosis and synaptic vesicle formation (4, 6-8) In this context, our findings bring a novel player into the glioma biology, which appears to have essential consequences for oligodendroglioma progression.

TRIM67 has been shown to have a dichotomous role in different cancers. It plays a tumor suppressive role in colon, gastric, lung and neuroblastoma tumors (10-12, 18, 19). In contrast, TRIM67 acts as an oncogene in non-small cell lung cancer (NSCLC)(13). Our study illustrates that oligodendroglioma is another example of a tumor in which TRIM67 acts as an oncogene. The wide range of mechanistic and phenotypic roles of TRIM67 in different tumors suggests that its function is highly tissue dependent. An interesting point to address in future studies could be to determine the common features between oligodendrogliomas and NSCLC that permit the oncogenic functions of TRIM67 to emerge.

Several members of the TRIM family have been associated with carcinogenesis and act to either promote or suppress tumor growth (3). In this context, TRIM67 emerges as another TRIM family member with the oncogenic function in glioma. The TRIM protein family also has E3 ubiquitin ligase activity, and direct ubiquitination targets of TRIM67 has so far been restricted to 80-K-H, such that TRIM67-induced degradation of 80-K-H leads to alteration of Ras signaling in neuroblastoma(10). A recent MS-based study detected potential TRIM67 and TRIM9 ubiquitination targets using mouse primary cortical neurons that need further functional characterization (20). In addition, our MS and co-IP-MS experiments using human glioma cell lines also provide datasets to further identify potential TRIM67 targets and interaction partners. It will be also of interest to compare TRIM67 interactome between human glioma and mouse cortical neurons to discover the key TRIM67 targets underlying tumorigenesis.

The crosstalk between TRIM9 and TRIM67 also requires further investigation. It has been suggested that TRIM9 and TRIM67 can act in antagonistic ways, regulating VASP degradation and filopodia extensions (6). However, ubiquitylome analysis in mouse cortical neurons indicated overlapping and non-overlapping targets of TRIM9 and TRIM67, suggesting that both E3 ligases could have redundant and non-redundant functions (20). Our study shows an enrichment of TRIM9 protein levels, physical interaction and colocalization of TRIM9 with TRIM67. However, further efforts are still required to comprehensively elucidate TRIM67/TRIM9 interaction axis as well as the crosstalk with other TRIMs in gliomagenesis.

Recent efforts have also focused on mapping the TRIM67 interactome. Using Bio-ID-MS approach, Menon *et. al*. reported that the interacting partners of TRIM67 majorly consist of cytoskeletal, synaptic, exocytosis and endocytosis proteins in mouse cortical neurons(8). Our co-IP-MS and total lysate-MS experiments using patient-derived glioma cell lines also demonstrate that TRIM67 interactome majorly contains cytoskeleton, endocytosis, cell adhesion and junction proteins. Thus, our study also to some extent detects similar pathways and components to Menon *et. al*. Moreover, our work also identified additional TRIM67-targeted pathways, such as Rho GTPase, RhoA and Rac signaling. These altered signaling pathways may underlie glioma pathogenesis. For example, metastatic cancer cells benefit from both elongated and rounded modes of migration(21). These two different modes of movement are regulated by Rho GTPases. Elongated cell migration is driven by RAC1 activity and coupled with F-actin-containing protrusions and extracellular matrix (ECM) degradation. On the other hand, bleb-based rounded cell migration is triggered by RHOA/C signaling-mediated ROCK activity and accompanied by actomyosin contractility without the necessity of ECM proteolysis (22). Bleb-based rounded cell motility prevails when active Rho-ROCK signaling is coupled with suppressed RAC1 signaling (15). Consistently, our findings also indicate activated Rho-ROCK but inhibited RAC1 signaling, which potentially underlies the TRIM67-driven membrane blebbing.

From a translational perspective, a comprehensive evaluation of transcriptional alterations that can distinguish between IDH mutant subtypes can ultimately either act as a therapeutic target or a pathological biomarker to distinguish subtypes when additional markers are needed. To this end, our work uncovered a functional pathogenic mechanism and laid the foundation for a novel therapeutic strategy to treat a disease that is now largely incurable. Rho-ROCK inhibition may also be considered as an additional or complementary therapy option for oligodendrogliomas. Additionally, targeting bleb-based cell motility might be an alternative strategy. Such therapeutic options could also be investigated in combination with recently developed clinical trial therapies for IDH mutant glioma, including IDH inhibitors (23) or IDH vaccine (24). Molecular and functional characterization of TRIM67 expression using patient-derived pathological samples could make a substantial progress in the brain tumor therapy and ultimately develop new strategies toward patient stratification and personalized medicine to improve the survival of glioma patients. In summary, these results highlight that the inhibition of TRIM67-mediated pathways as a viable therapeutic target in gliomas and the potential use of TRIM67 as a pathological biomarker unique to oligodendrogliomas.

## Materials and Methods

### Cell Culture

Human neural progenitor cells (hNPCs) and patient-derived glioma lines TS603, S24, TS600 and TS543 were maintained in Neurocult Basal Medium with proliferation supplements, 20 ng/ml EGF, 20 ng/ml basic-FGF and 2 μg/ml Heparin (StemCell Technologies). S24 Tdtomato/GFP cells were kindly gifted by Dr. Varun Venkataramani. HEK293TN and SK.N.BE2 cell lines were maintained in DMEM complete medium containing 10% FBS. Blebbistatin (Merck) treatment (20 μM) was performed for 1 h. Fasudil (HA-1077, Selleckchem) treatment (10 μM) was performed for 16 h. For the transient transfection of HEK cells, 300,000 cells/well were seeded on 6-well plate one day before to reach 70-80 % confluency on the day of transfection. 2 μg of total plasmid DNA/well were used in appropriate combinations of the plasmids: pLVX-puro (Addgene), pLVX-puro-TRIM67-Flag, pLenti-TRIM9-C-Myc-DDK-P2A-Puro (Origene), pLenti-TRIM9-C-mGFP (Origene). The reaction mix was filled up to 104 μl with OptiMEM (Gibco). Next, 6 μl of FuGene (Promega) transfection reagent was added. After 15 times gentle pipetting, the reaction was incubated at room temperature (RT) for 5-10 min. Cells were treated with the solution dropwise while gently shaking the plate and left in the incubator at 37 °C overnight. Similarly, CellLight Reagents BacMam 2.0 (Life Technologies) were also added to 70 % confluent HEK cells in a 2 μl/10,000 cells ratio and incubated for 16 h at 37 °C before imaging.

### Cloning of pLVX-puro-TRIM67-Flag overexpression plasmid

FLAG-TRIM67 was cloned into pLVX-puro (Addgene) using appropriate restriction enzyme sites. Digestion was performed at 37 °C for 4 h. Digests with 6x loading dye (NEB) were run on a 1 % agarose gel at 100 V for 1 h. Bands were extracted and cleaned up using QIAquick Gel Extraction Kit (Qiagen). Inserts were amplified in a thermal cycler. Insert amplification was performed using the Kappa Hi Fi Hotstart ReadyMix (Roche). For a 25 μL reaction, 12.5 μL of Kappa Hi Fi Hotstart ReadyMix (Roche) and each 0.75 μL (0.3 μM) of 10 μM forward and reverse primers was pipetted into a reaction tube. 70 ng of template DNA were added, and reaction volume adjusted to 25 μL using PCR-grade water. Amplification was followed by column clean ups using QIAquick PCR Purification kit (Qiagen). Amplified insert was each double-digested for 4 h at 37 °C, using restriction enzymes (NEB). Amplified inserts were run on gel and cleaned up using QIAquick Gel Extraction Kit (Qiagen). Ligations were performed using Quick Ligation Kit (NEB). Reaction mixes were incubated for 5 min at RT. Transformations of Stabl3 competent E-coli were performed using 5 μl of reaction mix per 50 μL competent bacteria. After confirming the positive clones using colony PCR and sanger sequencing, the plasmid was extracted using Maxiprep kit (Qiagen).

### Viral transductions

TRIM67 was stably overexpressed in hNPC, TS603, S24, TS600 and TS543 cell lines by viral transduction using pLVX-puro (Addgene), pLVX-puro-TRIM67-Flag plasmids. Additional luciferase expression was stably induced by a second viral transduction using pHIV-luc-ZsGreen (Addgene) plasmid and sorted by flow cytometry. SK.N.BE2 cells were infected with TRIM67 Mission lentiviral shRNA plasmids (Sigma). For virus production, HEK293TN cells were seeded in 6-well plates and transiently transfected with plasmid of choice using FuGene transfection reagent (Promega) and OptiMem (Gibco). Cells were transfected with packaging psPAX2 (Sandrine Sander), envelope pCMV-VSV-G (Addgene) plasmid and expression plasmid in a ratio of 1:1:1 ratio for virus production. Cell culture medium was changed after 24 hours and collected for harvesting at 24 and 48 hours after transfection. After the first and second harvest, the cell culture supernatant was mixed in 3:1 ratio to a lab-made filtered 4x virus precipitation buffer (40 g PEG 8000, 7g NaCl, 10 ml PBS, final pH adjusted to 7.0 with 1N HCl, filled up to 100 ml with distilled water) and incubated overnight at 4 °C. Next, the PEG/medium mix was centrifuged at 1600 x g at 4 °C for 45 min for virus collection. Supernatant was discarded, virus pellets were resuspended in PBS and added to the cell lines. To select transduced cells, 0.5 μg/mL puromycin was added every three days until cells are stable.

### SDS-page and western blot

Cells were directly lysed in the wells using appropriate volume of M-PER mammalian protein extraction reagent (Thermofisher) containing 1x Protease Inhibitor Cocktail (Roche). Lysates were sonicated with Sonorex Digitec (Bandelin) for 5 min, centrifuged at 16,000 g for 10 min at 4 °C and the collected in 1.5 ml microcentrifuge tubes. Lysates were then either used for BCA Protein Assay according to manufacturer’s protocol (ThermoFisher) or supplemented with 4X Laemmli Buffer (BioRad) containing 10 % β-mercaptoethanol and boiled at 95 °C for 5 minutes followed by Western Blot. 10 % SDS-PAGE gels were used for the electrophoresis. Equal amounts of protein, defined by BCA assay or defined by cell seeding numbers were loaded on 10% gels. Blotting was performed to the nitrocellulose membrane (Bio-Rad) using the Mini-PROTEAN Tetra electrophoresis wet blot system (Bio-Rad) at 100 V for 120 min. Membranes were blocked for 30 min in 5 % milk. Primary antibodies were applied overnight at 4 °C in 5% milk. Protein signal was detected using Pierce ECL Western Blotting Substrate (Thermo Fisher) and ChemiDoc XRS+ Gel Imaging System (Bio-Rad). If further probing on same membranes was performed, membranes were reconstituted using Restore PLUS Western Blot-Stripping-Buffer (ThermoFisher) for 2 times 15 minutes, followed by a TBST washing and incubation of membranes in 5 % milk before adding the next primary antibody ON at 4 °C. For GTP-pulldown experiments, TS603 cells were washed and seeded in neural stem cell media without EGF/bFGF supplementation. Cells were kept serum starved for 24 h, afterwards medium was changed to neural stem cell media containing 50 ng/μl EGF. Cells were then collected after 15 min EGF induction and flash frozen in liquid nitrogen. The following lysate preparation, GTP-pulldown and western blot procedures were performed according to RHOA/RAC1/CDC42 Activation Assay Combo kit (Cytoskeleton) manufacturer’s protocol. Primary antibodies used were as follows: mouse anti-RAC1 (Cytoskeleton #ARC03, 1:250); mouse anti-RHOA (Cytoskeleton #ARH05, 1:500); rabbit anti-ROCK2 (Cell Signaling #9029, 1:1000); rabbit anti-Tensin-2 (Cell Signaling #11990, 1:1000); rabbit anti-FAK (Cell Signaling #3285, 1:1000); rabbit anti-phospho-FAK (Cell Signaling #8556, 1:1000); mouse anti-Flag (Sigma #F1804, 1:1000); mouse anti-Akt (Cell Signaling #2920, 1:2000); rabbit anti-phospho-Akt (Cell Signaling #4060, 1:2000); rabbit anti-Vimentin (Cell Signaling #3932, 1:1000); rabbit anti-GAPDH (Cell Signaling #2118, 1:1000); rabbit anti-MAP1B (Merck #MAB366, 1:500); mouse anti-TRIM9 (Abnova #ABN-H00114088-M01, 1:1000); rabbit anti-GFP (Novus Bio #NB600-308, 1:1000). Secondary antibodies used were as follows: HRP-linked anti-mouse IgG (Cell Signaling #7076, 1:5000); HRP-linked anti-rabbit IgG (Cell Signaling #7074, 1:5000).

### Orthotopic transplantation

All mouse experiments were approved by the Institutional Animal Care and Use Committee at DKFZ. Female athymic nude mice at age of 7 weeks (n=15 per CTRL/TRIM67 groups; n= 16 for IVIS imaging, n=14 for survival) were intracranially injected using a fixed stereotactic apparatus (Stoelting). Mice were weighed and injected with 5 mg/kg Carprofen subcutaneously. Mice were then put into an induction chamber with 1-3 Vol% Isoflurane. Once mice did not show reflex to toe-pinching, mice were put into stereotactic frame, on a heat mat on 36 °C, isoflurane was provided during and following surgery via nose cone using 1-2 Vol% Isoflurane. Mouse eyes were covered with eye cream to prevent dryness. Mice were injected with 2 mg/kg bupivacaine near incision site for additional local anesthesia. Iodine solution was used on the mouse head as a disinfectant for incision site. A vertical cut was made to expose the skull and a small hole was drilled oriented 1.5 mm above and 1.5 mm right of Bregma. Hamilton syringe was filled with 25,000 cells/μL cell solution. Injection of cells was performed using the previously drilled hole to access brain, syringe tip was positioned using stereotactic frame. 50,000 cells were injected 1.5 mm deep into the cortex at a speed of 1 μL/min. Cells were allowed to settle for 5 min before syringe was removed slowly. Incision site was closed using tissue glue.

### Bioluminescence imaging

Cell lines for use in orthotopic in vivo experiments were labeled with pHIV-luc-ZsGreen (Addgene). Bioluminescent imaging was performed every 3-7 days following intraperitoneal injection of D-luciferin and measured using the Xenogen IVIS Spectrum in vivo imaging system (PerkinElmer). Living Image software (PerkinElmer) was used to acquire and analyze the BLI data.

### Immunofluorescence

Cells were seeded on matrigel in a concentration of 50,000 cells/well in 24-well dishes in 500 μL of medium. For the staining, 24 h after seeding, medium was removed, and cells were washed twice using PBS. 500 μL 4 % PFA was added to each well and incubated for 20 min at room temperature. PFA was removed and cells were washed twice with PBS. Cells were permeabilized using PBS-T (containing 0.1 % of Triton X-100) for 10 min at room temperature. Cells were blocked using 5% BSA/PBS-T buffer for 30 min at RT. Primary antibody was diluted in 5% BSA/PBS-T buffer and cells were incubated at 4°C for overnight. Primary antibody was removed, and cells were washed three times using PBS-T. Secondary antibody was added to the cells in 5% BSA/PBS-T buffer and incubated for 1 h at room temperature. Afterwards, cells were washed twice using PBS-T, incubated in DAPI for 5 min and washed once with PBS. Coverslips were mounted to microscopy slides using 1 drop of mounting media (VECTASHIELD) per coverslip. Primary antibodies used for IF staining were: Phalloidin-iFluor 594 conjugated (Abcam #176757, 1:1000); mouse anti-Flag (Sigma #F1804, 1:200); mouse anti-TRIM9 (Abnova #ABN-H00114088-M01, 1:200); rabbit anti-MAP1B (Merck #MAB366, 1:200); rabbit anti-cleaved-CASP3 (Cell Signaling #9664, 1:500); rabbit anti-cleaved-PARP (Cell Signaling #5625, 1:400). Secondary antibodies used for IF staining were: Alexa Fluor 488 goat anti-mouse (Thermofisher #A-11029, 1:1000); Alexa Fluor 488 donkey anti-rabbit (Thermofisher #A-21206, 1:1000); Alexa Fluor 594 goat anti-mouse (Thermofisher #A-11032, 1:1000); Alexa Fluor 594 donkey anti-rabbit (Thermofisher #A-21207, 1:1000).

### Microscopy imaging and quantification

The image z-stacks acquisition of DAPI, Alexa-488 and Alexa-594 immunofluorescent-stained cells was performed with Leica TCS SP5 II Confocal microscope equipped with UV 405nm diode, Argon and HeNe 594nm lasers, using 63x / 1.40 PL APO objective and PMT detectors, all controlled by LAS AF software. The 3D z-stack projections were prepared using Fiji version of ImageJ (25). F-Actin intensity quantification was performed using in-house developed FIJI macros. Image acquisition for IHC-stained mice brain sections were performed using ZEN software on Zeiss Cell Observer. Z1 microscope in DKFZ light microscopy facility. The microscope setup for bright field imaging, Koehler illumination, tile region adjustment and stitching were performed according to SOPs provided by the facility.

### Embedding, microtome sectioning, immunohistochemistry and H&E staining

When the endpoints were reached, animals were euthanized, and the mice brains were dissected out. The brains were then formalin-fixed, paraffin-embedded (FFPE) and microtome-sectioned until the tumor injection site (1.5 mm before bregma) by Gabriele Schmidt according to the full project routine SOPs of DKFZ histology facility. For IHC and H&E staining, the FFPE tissue sections were selected from two specific regions of the brain: 1) Early region (roughly around 2.25 mm before bregma), 2) Late region (roughly around 1.5 mm before bregma), and staining protocols were performed by Claudia Schmidt. For Flag-IHC staining, 5-micron thick sections were deparaffinized, rehydrated, microwave-pretreated with citrate buffer pH 6.0 for 20 min, cooled-down to RT and air-dried. After framing with Dako-Pen and washing with TBST, tissue sections were blocked with 3% H2O2 in TBST for 10 min at RT and followed with two times TBST wash. Then, the following treatments were sequentially applied: treatment with 10% donkey serum/TBS for 20 min at RT, 2x wash with TBST, Avidin treatment for 10 min at RT, 2x wash with TBST, Biotin treatment (Avidin/Biotin Blocking Kit, Vector # SP-2001) for 10 min at RT and 2x wash with TBST. Rabbit monoclonal anti-Flag (abcam # ab 205606, 1:500) was applied in DAKO antibody diluent (DAKO # S2022) with 2% dry milk for 2 h at 37°C and followed by 3x wash with TBST. Biotinylated goat anti-rabbit IgG (dianova # 111-065-144, 1:400) in TBST was applied for 30 min at 37°C and 3x washed with TBST. Next, Alkaline Phosphatase Streptavidin (Vector # SA-5100, 1:400) in TBST was added for 30 min at 37°C. Substrate RED (DAKO Kit # K5005) was then used for 10 min at RT and rinsed in distilled water. It was followed by haemalaun treatment for 1 min, rinse with tap water for 10 min and mounting with Aquatex (Merk # 108562). For Ki-67 IHC staining, similar protocol was performed with only few differences. In this case, sections were pretreated with steamer using Tris buffer pH 9,0 for 20 min. The blocking was done with 2% horse serum. Rabbit monoclonal anti (human) Ki-67 (abcam # ab 16667, 1:100) was performed as primary antibody.

H&E staining was performed applying treatments in the following order: 1) Xylol for 10 min twice, 2) ethanol for 5 min twice, 96% Alcohol for 5 min, 3) 80 % alcohol for 5 min, 4) 70 % alcohol for 5 min, 5) aqua dest for three times, 6) haemalaun (Roth # T865.3) for 2 min, 7) rinse with tap water for 5 min, 8) eosin (Merck. #1.15935.0100) for 1 min, 9) wash with aqua dest, 10) dehydration up to 100% ethanol, 11) Xylol twice, 12) mounting with Eukitt.

### Wound healing

SK.N.BE2 and TS600 lines were seeded in a density of 5×10^5^ cells/well on 6-well plates, and they were incubated for 3-5 days until they reach 100% confluency. The wound was simultaneously created using a P100 pipette tip in the middle of each well. The images were taken after 24 h and 48 h using Nikon Eclipse Ts2 bright field microscope at 4x magnification. The wound area was quantified using in-house developed FIJI macros.

### Cell adhesion

48-well ECM cell adhesion assay (Cell Biolabs) was performed based on the manufacturer’s protocol. Briefly, TS603 CTRL and TRIM67 cells were seeded in a density of 5×10^4^ cells/well on a 48-well ECM-coated plate, and they were incubated for 2.5 hours in EGF/FGF-free medium. After discarding the media, wells were washed 4 times with 250 μl PBS. Then, 200 μl cell stain solution was added and incubated for 10 min at RT. Wells were gently washed 4 times with 500 μl deionized water and air-dried. The image of the plate was taken by ChemiDoc XRS+ imaging system (Biorad). 200 μl extraction solution was added and incubated for 10 min on an orbital shaker. 150 μl of each extracted sample was transferred to 96-well plate, and OD was measured at 560 nm in Infinite 200 Pro plate reader (Tecan).

### Bioinformatic analysis

RNA-seq expression for low-grade glioma were obtained from The Cancer Genome Atlas (https://www.cancer.gov/tcga), and ATAC-seq data were obtained from Corces *et al* (26). Normalized RNA-seq expressions were used from low-grade gliomas in TCGA. EPIC methylation array data obtained from the Department of Clinical Pathology at Heidelberg University Hospital as described previously (27). The methylation data was analyzed with R3.5.3/minfi 1.28.4 (28) and limma package (29). Beta values from EPIC methylation arrays included two relevant probes at TRIM67 promoter region. Two open chromatin peaks were also identified by ATAC-seq and the score value from the paper was used.

### RNA-sequencing

hNPC, TS603, S24, TS600 and TS543 cells were seeded in a concentration of 5 × 10^5^ cells/well in 6-well dishes in triplicates and harvested 48 h after seeding. Total mRNA from biological triplicates of CTRL/TRIM67was extracted using the QIAGEN RNeasy RNA isolation kit according to manufacturer’s protocol. Extracted RNA was tested for quality using the Bioanalyzer system by Agilent following manufacturer’s instructions. The sequencing of samples was performed by the DKFZ genomics and proteomics core facility. RNA quantity and quality were validated using 4200 TapeStation (Agilent), and samples with above an RNA integrity number of 9.8 were used for sequencing. RNA-seq libraries were created with TruSeq Stranded mRNA Library Prep Kit (Illumina). The sequencing performed was HiSeq 4000 Single-read 50 bp. The aligned bam files and feature counts were generated by the Omics IT and data management core facility at DKFZ (ODCF). Normalization of the raw counts and differential expression analysis were performed using DESeq2 R package (30) and Galaxy (31). An adjusted p-value of 0.05 was used as a cutoff to stringently filter the significantly altered genes. Volcano plots and heatmaps were created using Graphpad Prism 9. Enriched pathways were determined using the Ingenuity Pathway Analysis (QIAGEN Inc., https://www.qiagenbioinformatics.com/products/ingenuity-pathway-analysis) (32) or Enrichr (33-35).

### Immunoprecipitation and mass spectrometry

Whole cell lysates from duplicates per condition, each 5 × 10^6^ cells, were prepared using the IP buffer (50 mM Tris pH 7.5, 150 mM NaCl, 1% TritonX-100, 0.5% Na-DOC, 1 mM EDTA, 2 mM PMSF and 1x Roche protease inhibitor cocktail). After sonification for 5 min and clarification with centrifuge at 16,000 g at 4 °C, 2 μg of Flag (Sigma #F1804) antibody were added to each CTRL/TRIM67 IP samples. The reaction volumes were adjusted to 1 ml using IP buffer. Reaction was incubated for 1 h at 4 °C on a rotator. 25 μL of Protein G agarose beads (Roche) were placed into each IP sample and incubated o/n at 4 °C on a rotator. After incubation, bead-antibody-protein conjugates were washed in 1 mL of IP/Wash Buffer for 6 times. Next, pulled-down proteins were eluted from the beads using 2x laemmli buffer (Bio-Rad) at 95 °C for 5 min incubation in heat block at 1000 rpm. Proteomics were performed at the Mass Spectrometry Core Facility at the German Cancer Center (DKFZ).

### Statistical analysis

Two-sided t-test was performed if not stated otherwise using Prism 9. Significance is indicated using following legend: ≤0. 05 (*), ≤0.01 (**), ≤0.001 (***). Survival analysis was also performed using log-rank test in Prism 9.

## Accession Numbers

All the RNA-seq data have been deposited in NCBI’s Gene Expression Omnibus under accession number GSE184567.

## Acknowledgements

We thank the members of the Turcan lab for helpful discussions. We thank the Genomics and Proteomics Core Facility (GPCF) at the DKFZ for providing next-generation sequencing (NGS) services and proteomics services and analysis. We thank the Omics IT and Data Management Core Facility (ODCF) at the DKFZ for data management and technical support. We thank Dr. Varun Venkataramani for providing us S24 Tdtomato/GFP cells.

This work was supported by the German Cancer Aid, Max Eder Program grant number 70111964 (S.T.) and DFG Project-ID 404521405, SFB 1389 – UNITE Glioblastoma, Work package A04 (S.T.). J.T. is supported by a DFG Mercator Fellowship (DFG grant number TU 585/1-1).

## Author Contributions

E.D. and S.T. designed and directed the study. R. A. identified TRIM67 upregulation from bioinformatic analyses. E.D., R.A., A.N., X.S., J.P.V. and B.S. performed the experiments. E.D., R.A., A.N. and S.T. analyzed the data. E.D., A.N., J.T. and S.T interpreted the data. M.B., D.K., C.S. and G.S. provided technical assistance. D.K. provided custom-built macro plugins for image analysis. F.B. and J.T. provided conceptual advice. E.D. and S.T. wrote the paper. All authors contributed to the writing and/or editing of the manuscript.

---

**Figure S1**. TRIM67 overexpression influences cytoskeletal pathways and proteins. **A)** The bar graph comparison of log_2_ expression for *TRIM67* between different tumor types using TCGA data. **B)** Immunoblotting for Flag-TRIM67 and actin under control and TRIM67 overexpression conditions in hNPC, S24, TS603 cells (upper panel); TS600 and TS543 cells (lower panel). Actin serves as a loading control. **C)** Volcano plot for TS603_MS. The significance threshold for -log_10_(padj) was set to 1.4. Thresholds for log_2_ fold change are -1 and 1. The proteins above and below log_2_ fold change thresholds are depicted in red, and the rest of the proteins are depicted in gray. The proteins of interest are marked and shown on the plot. **D)** Volcano plot for S24_MS. The significance threshold for -log_10_(p adjusted value) was set to 1.4. Thresholds for log_2_ fold change are -1 and 1. The proteins above and below log_2_ fold change thresholds are depicted in red, and the rest of the proteins are depicted in gray. The proteins of interest are marked and shown on the plot. **E)** Top 10 significantly-enriched cytoskeletal pathways from TS603_MS experiment. The significance threshold for -log_10_(p value) was set to 1.4, which is shown by the dashed red line. **F)** Top 10 significantly-enriched cytoskeletal pathways from S24_MS experiment. The significance threshold for -log_10_(p value) was set to 1.4, which is shown by the dashed red line. **G)** Heatmap comparison of actin cytoskeleton signaling pathway for S24_MS, Co-IP_MS and TS603_MS experiments using IPA and Prism. **H)** The scheme of actin cytoskeleton signaling created by using IPA. The proteins marked in red are upregulated, whereas those marked in green are downregulated in S24_MS experiment.

**Figure S2**. TRIM67 partially colocalizes with and influences the cytoskeletal proteins. **A)** Representative IF images of HEK293-TN cells upon control and TRIM67 overexpression conditions. DIC indicates bright field images, Flag-TRIM67 in green, ActRFP in red and DAPI in blue. The cells of interest are marked by white triangles. Scale bar indicates 25 μm. **B)** Representative IF images of HEK293-TN cells upon control and TRIM67 overexpression conditions. DIC indicates bright field images, Flag-TRIM67 in green, TubRFP in red and DAPI in blue. The cells of interest are marked by white triangles. Scale bar indicates 25 μm. **C)** Representative IF images of HEK293-TN cells upon control and TRIM67 overexpression conditions. DIC indicates bright field images, Flag-TRIM67 in green, MAP1B in red and DAPI in blue. The cells of interest are marked by white triangles. Scale bar indicates 25 μm. **D)** Immunoblotting for MAP1B and GAPDH under control and TRIM67 overexpression conditions in S24 and TS603 cells. GAPDH serves as a loading control.

**Figure S3**. TRIM67 interacts with TRIM9 and changes its expression. **A)** Immunoblotting for TRIM9 and GAPDH under control and TRIM67 overexpression conditions in S24, TS603 and TS600 cells. GAPDH serves as a loading control. **B)** The normalized quantification of part A for TRIM9. Error bars indicate the standard error of the mean. Two-sided t-test was performed, * p<0.05 and ** p<0.01. **C)** Immunoblotting for TRIM9 and Flag-TRIM67 following α-Flag pulldown under control and TRIM67 overexpression conditions in S24 cells. **D)** Immunoblotting for TRIM9-GFP and Flag-TRIM67 following α-Flag pulldown under control and TRIM67 overexpression conditions in HEK 293-TN cells. **E)** Representative IF images of HEK293-TN cells co-transfected with either control/TRIM9-GFP or Flag-TRIM67/TRIM9-GFP. GFP in green, Flag in red and DAPI in blue. Scale bar indicates 25 μm. **F)** Representative IF images of HEK293-TN cells co-transfected with either control/Myc-TRIM9 or Flag-TRIM67/Myc-TRIM9. TRIM9 in green, F-Actin in red and DAPI in blue. Scale bar indicates 10 μm.

**Figure S4**. TRIM67-overexpressing cells do not tend to be apoptotic. **A)** Representative IF images of TS600 cells upon control and TRIM67 overexpression conditions. Cleaved CASP3 is shown on the left panel, cleaved PARP is shown on the right panel. DIC indicates bright field images, cleaved CASP3/cleaved PARP in green, Flag-TRIM67 in red and DAPI in blue. CHX (20 ng/μl, 6 hours) treatment, as an apoptotic inducer, was used for positive control. **B)** Representative IF images of TS603 cells upon control and TRIM67 overexpression conditions. Cleaved CASP3 is shown on the left panel, cleaved PARP is shown on the right panel. DIC indicates bright field images, cleaved CASP3/cleaved PARP in green, Flag-TRIM67 in red and DAPI in blue. CHX (20 ng/μl, 6 hours) treatment, as an apoptotic inducer, was used for positive control.

**Figure S5**. Cell migration and tumor growth is more pronounced *in vitro* and *in vivo* upon TRIM67 overexpression. **A)** Representative bright field images of the wound healing experiment at 0 h, 24 h and 48 h. Left panel shows SK.N.BE2 cells upon scrambled shRNA, TRIM67 shRNA#4 and TRIM67 shRNA#5 treatment. Right panel shows TS600 cells upon control and TRIM67 overexpression conditions. Scale bar indicates 200 μm. **B)** Bioluminescence images of mice on day 18 with orthotopic injection of TS603-CTRL and TS603-TRIM67 cells. The flux measurement of the tumor signals and empty regions for normalization is shown on the pictures. One early-euthanized (sac’d) TRIM67 animal is not shown here.

## References

1. D. J. Brat et al., Comprehensive, Integrative Genomic Analysis of Diffuse Lower-Grade Gliomas. N Engl J Med 372, 2481–2498 (2015).

2. H. Yan et al., IDH1 and IDH2 Mutations in Gliomas. New England Journal of Medicine 360, 765–773 (2009).

3. S. Hatakeyama, TRIM Family Proteins: Roles in Autophagy, Immunity, and Carcinogenesis. Trends Biochem Sci 42, 297–311 (2017).

4. N. P. Boyer, C. Monkiewicz, S. Menon, S. S. Moy, S. L. Gupton, Mammalian TRIM67 Functions in Brain Development and Behavior. eNeuro 5 (2018).

5. D. J. Montell, TRIMing Neural Connections with Ubiquitin. Dev Cell 48, 5–6 (2019).

6. N. P. Boyer, L. E. McCormick, S. Menon, F. L. Urbina, S. L. Gupton, A pair of E3 ubiquitin ligases compete to regulate filopodial dynamics and axon guidance. J Cell Biol 219 (2020).

7. F. L. Urbina et al., TRIM67 regulates exocytic mode and neuronal morphogenesis via SNAP47. Cell Rep 34, 108743 (2021).

8. S. Menon et al., The TRIM9/TRIM67 neuronal interactome reveals novel activators of morphogenesis. Mol Biol Cell 32, 314–330 (2021).

9. L. D. Do et al., TRIM9 and TRIM67 Are New Targets in Paraneoplastic Cerebellar Degeneration. Cerebellum 18, 245–254 (2019).

10. H. Yaguchi et al., TRIM67 protein negatively regulates Ras activity through degradation of 80K-H and induces neuritogenesis. J Biol Chem 287, 12050–12059 (2012).

11. S. Wang et al., TRIM67 Activates p53 to Suppress Colorectal Cancer Initiation and Progression. Cancer Res 79, 4086–4098 (2019).

12. R. Liu et al., TRIM67 promotes NFkappaB pathway and cell apoptosis in GA13315treated lung cancer cells. Mol Med Rep 20, 2936–2944 (2019).

13. J. Jiang et al., TRIM67 Promotes the Proliferation, Migration, and Invasion of Non-Small-Cell Lung Cancer by Positively Regulating the Notch Pathway. J Cancer 11, 1240–1249 (2020).

14. N. S. Clayton, A. J. Ridley, Targeting Rho GTPase Signaling Networks in Cancer. Front Cell Dev Biol 8, 222 (2020).

15. V. Sanz-Moreno et al., Rac activation and inactivation control plasticity of tumor cell movement. Cell 135, 510–523 (2008).

16. O. T. Fackler, R. Grosse, Cell motility through plasma membrane blebbing. J Cell Biol 181, 879–884 (2008).

17. S. Hafizi, F. Ibraimi, B. Dahlback, C1-TEN is a negative regulator of the Akt/PKB signal transduction pathway and inhibits cell survival, proliferation, and migration. FASEB J 19, 971–973 (2005).

18. W. Jin, J. Shi, M. Liu, Overexpression of miR-671-5p indicates a poor prognosis in colon cancer and accelerates proliferation, migration, and invasion of colon cancer cells. Onco Targets Ther 12, 6865–6873 (2019).

19. Y. Liu et al., TRIM67 inhibits tumor proliferation and metastasis by mediating MAPK11 in Colorectal Cancer. J Cancer 11, 6025–6037 (2020).

20. S. Menon, D. Goldfarb, E. M. Cousins, M. B. Major, S. L. Gupton, The ubiquitylome of developing cortical neurons. MicroPubl Biol 2020 (2020).

21. K. M. Yamada, M. Sixt, Mechanisms of 3D cell migration. Nat Rev Mol Cell Biol 20, 738–752 (2019).

22. J. B. Wyckoff, S. E. Pinner, S. Gschmeissner, J. S. Condeelis, E. Sahai, ROCK- and myosin-dependent matrix deformation enables protease-independent tumor-cell invasion in vivo. Curr Biol 16, 1515–1523 (2006).

23. L. Gatto et al., IDH Inhibitors and Beyond: The Cornerstone of Targeted Glioma Treatment. Mol Diagn Ther 25, 457–473 (2021).

24. M. Platten, L. Bunse, W. Wick, Emerging targets for anticancer vaccination: IDH. ESMO Open 6, 100214 (2021).

25. J. Schindelin et al., Fiji: an open-source platform for biological-image analysis. Nat Methods 9, 676–682 (2012).

26. M. R. Corces et al., The chromatin accessibility landscape of primary human cancers. Science 362 (2018).

27. R. Al-Ali et al., Single-nucleus chromatin accessibility reveals intratumoral epigenetic heterogeneity in IDH1 mutant gliomas. Acta Neuropathol Commun 7, 201 (2019).

28. M. J. Aryee et al., Minfi: a flexible and comprehensive Bioconductor package for the analysis of Infinium DNA methylation microarrays. Bioinformatics 30, 1363–1369 (2014).

29. M. E. Ritchie et al., limma powers differential expression analyses for RNA-sequencing and microarray studies. Nucleic Acids Res 43, e47 (2015).

30. M. I. Love, W. Huber, S. Anders, Moderated estimation of fold change and dispersion for RNA-seq data with DESeq2. Genome Biol 15, 550 (2014).

31. E. Afgan et al., The Galaxy platform for accessible, reproducible and collaborative biomedical analyses: 2018 update. Nucleic Acids Res 46, W537–W544 (2018).

32. A. Kramer, J. Green, J. Pollard, Jr., S. Tugendreich, Causal analysis approaches in Ingenuity Pathway Analysis. Bioinformatics 30, 523–530 (2014).

33. E. Y. Chen et al., Enrichr: interactive and collaborative HTML5 gene list enrichment analysis tool. BMC Bioinformatics 14, 128 (2013).

34. M. V. Kuleshov et al., Enrichr: a comprehensive gene set enrichment analysis web server 2016 update. Nucleic Acids Res 44, W90–97 (2016).

35. Z. Xie et al., Gene Set Knowledge Discovery with Enrichr. Curr Protoc 1, e90 (2021).

