## Supplementary figures and images for "TRIM67 drives tumorigenesis in oligodendrogliomas through Rho GTPase-dependent membrane blebbing"

### Supplemental Fig. 1

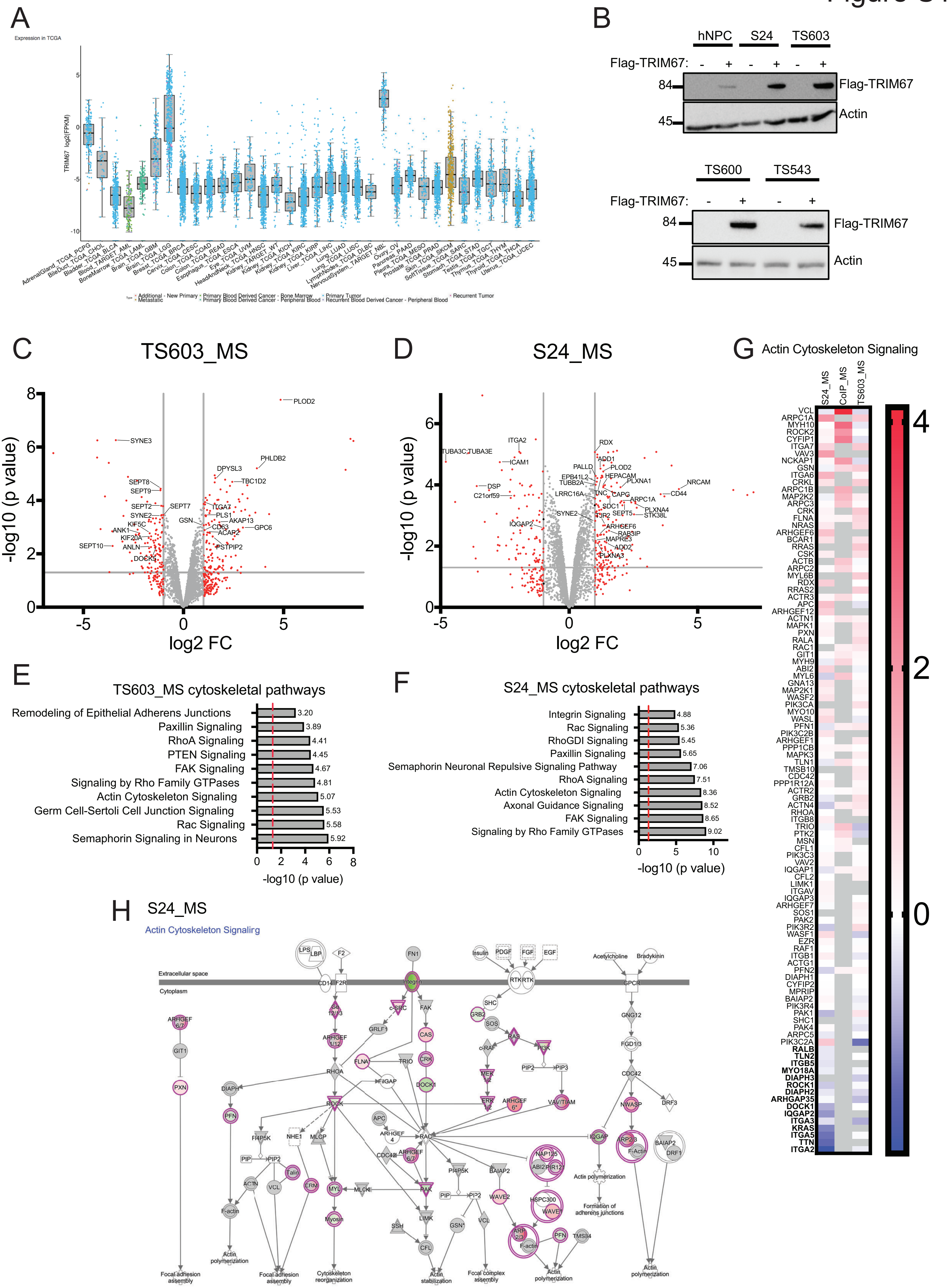

### Supplemental Fig. 2

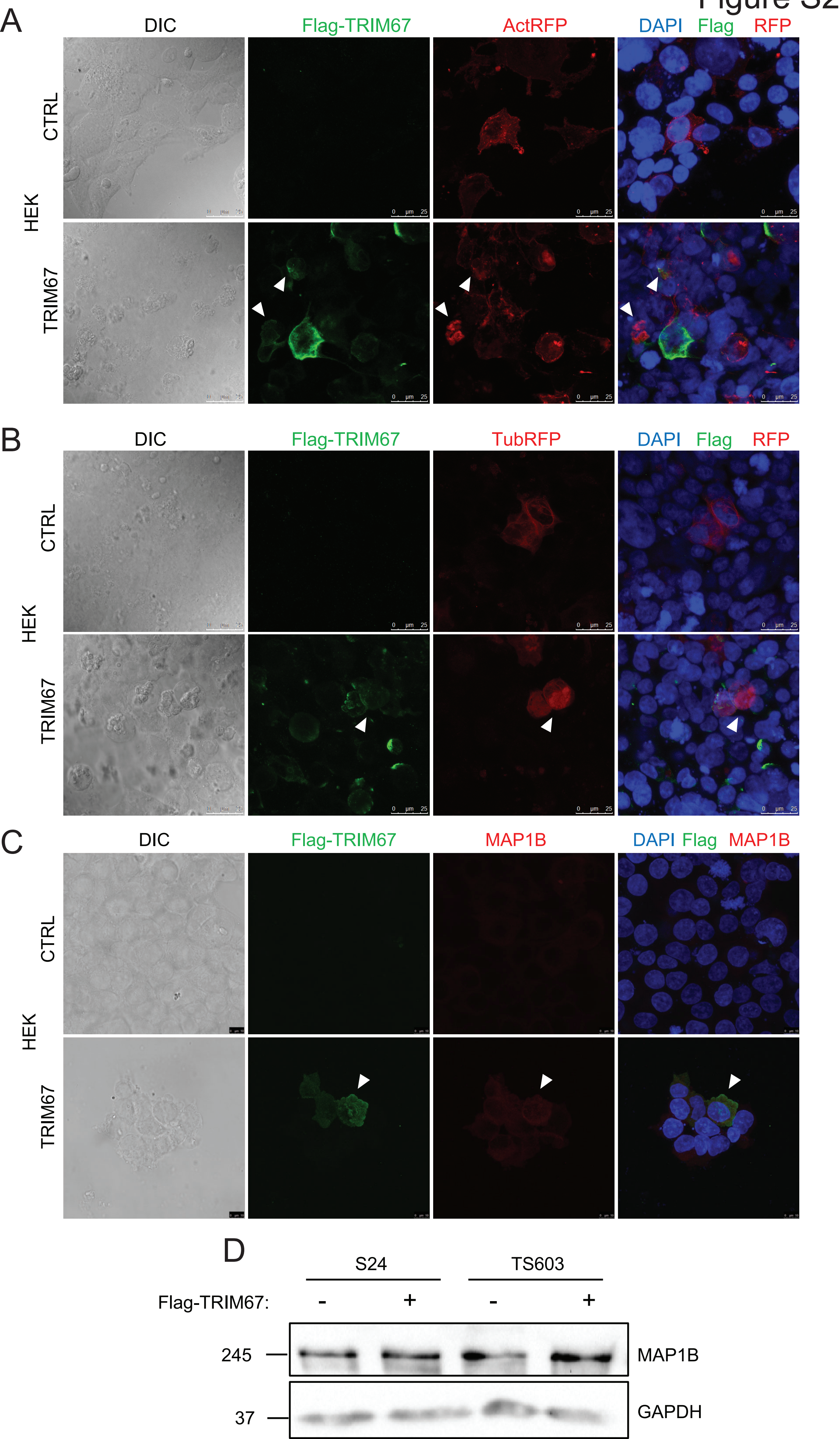

### Supplemental Fig. 3

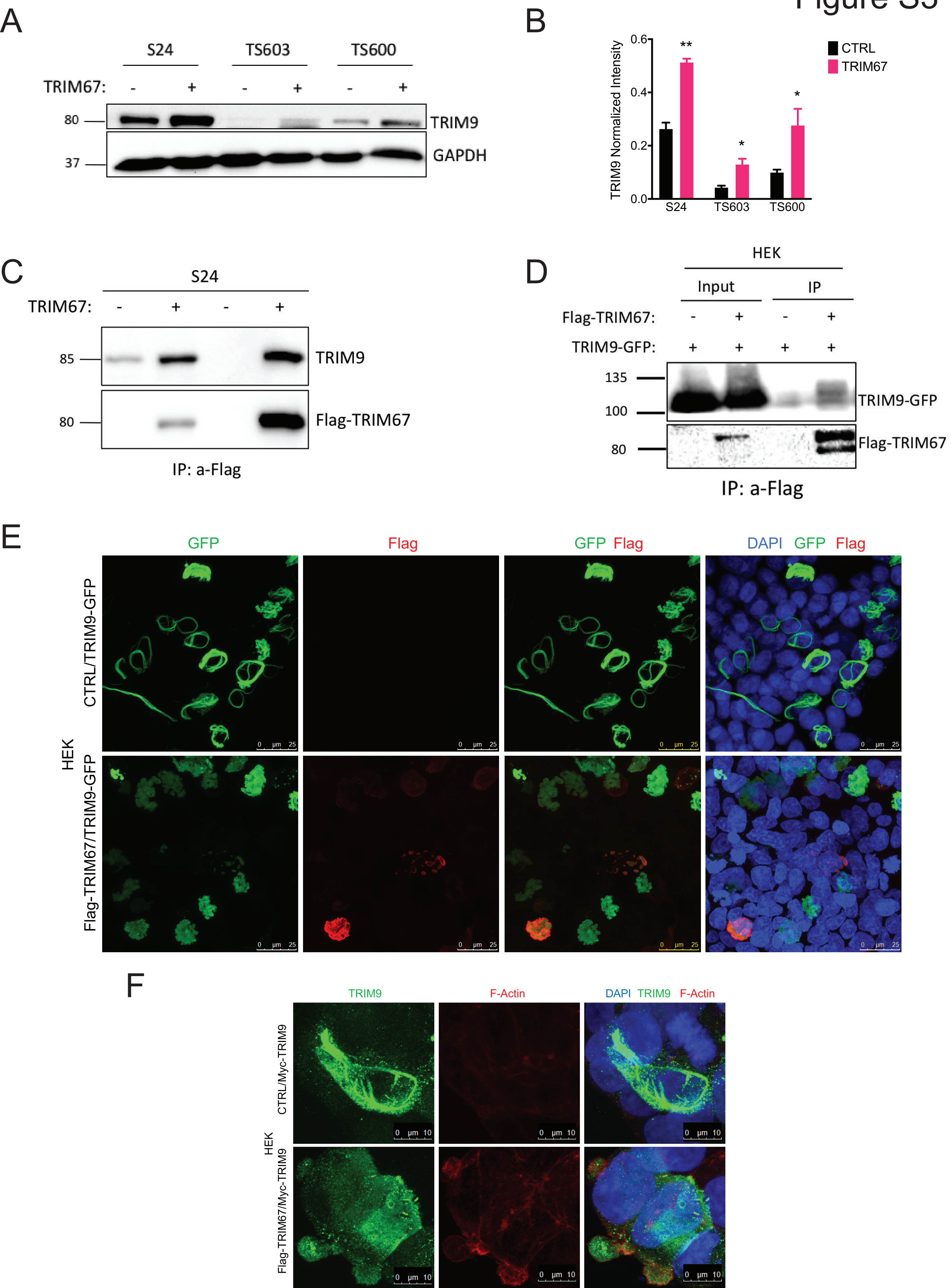

### Supplemental Fig. 4

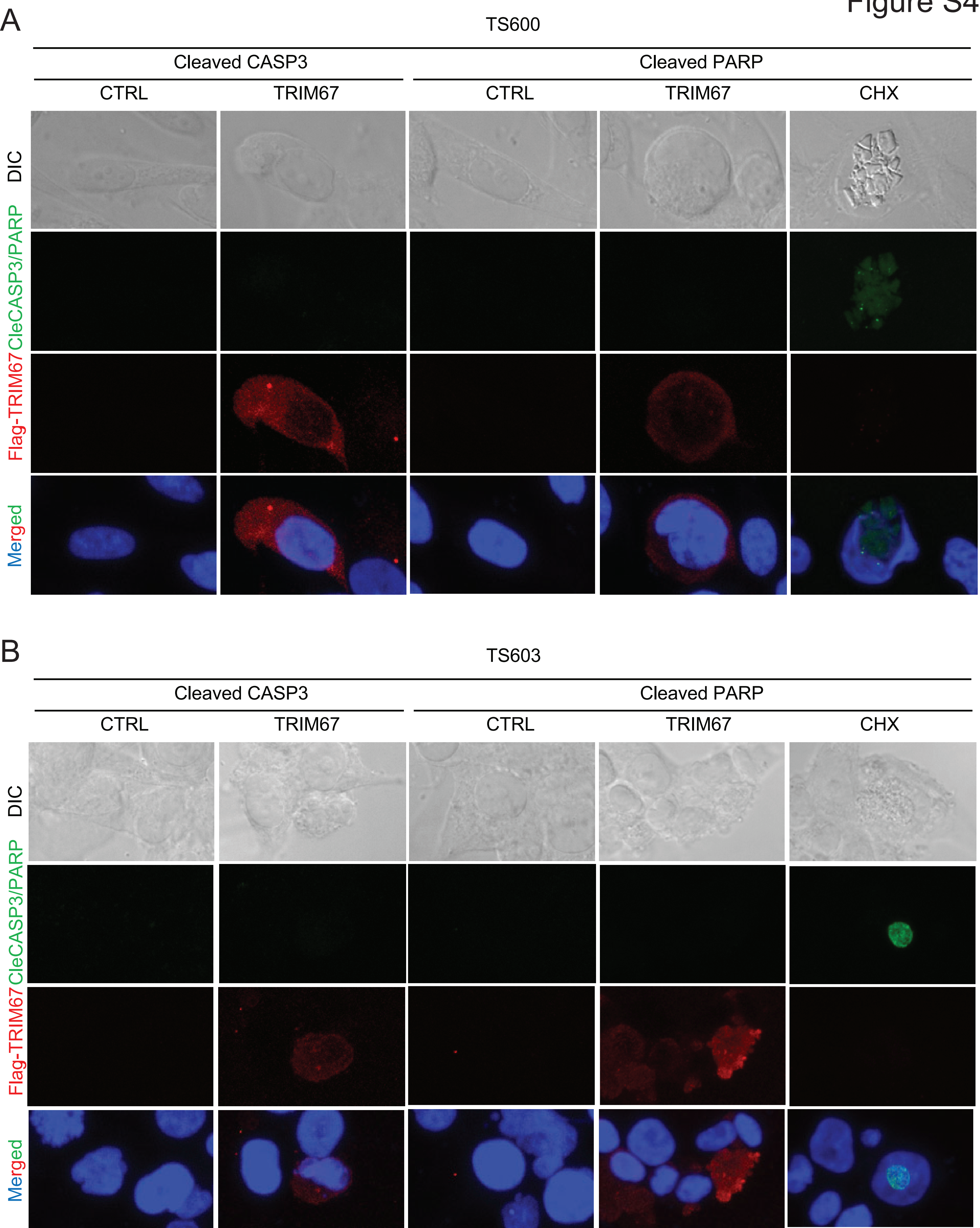

### Supplemental Fig. 5

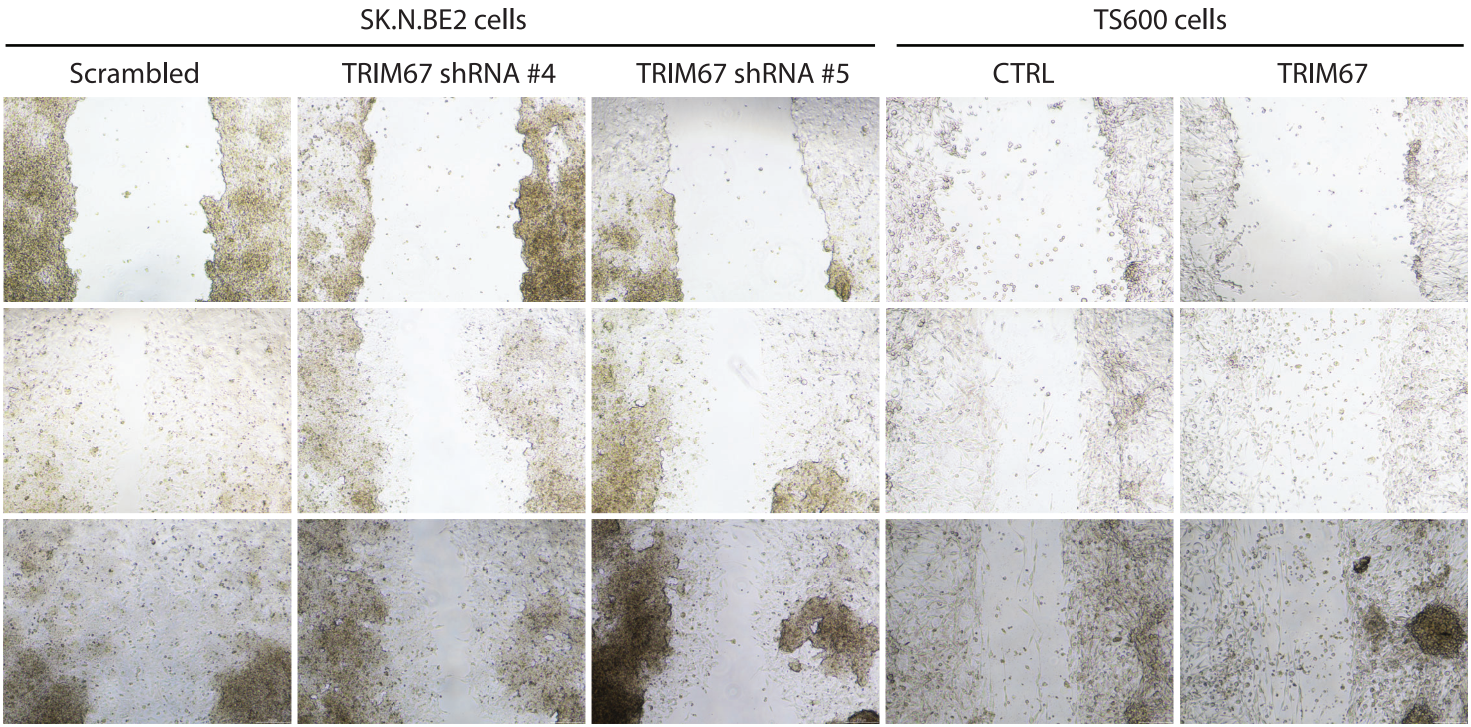

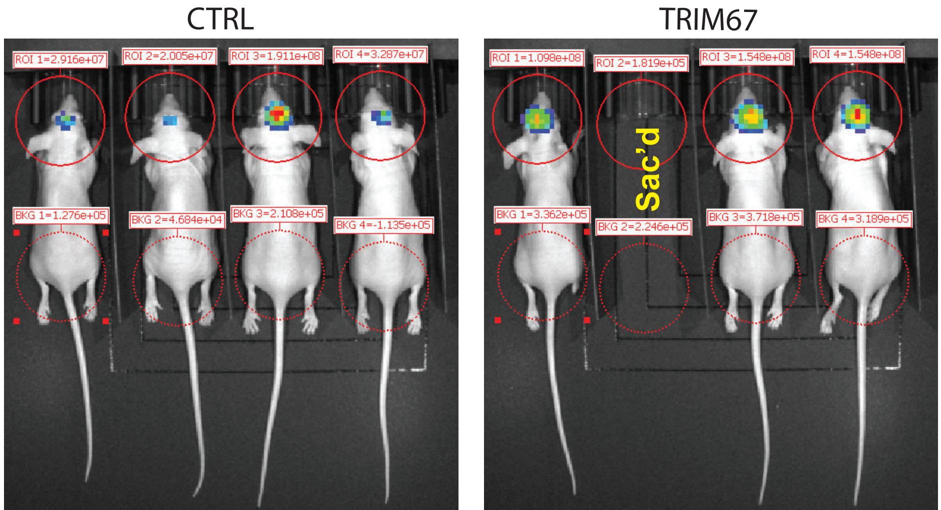
